# Effective macrophage clearance of *Klebsiella pneumoniae* requires the inducible nitric oxide synthase iNOS and is independent of reactive oxygen species generated by NADPH oxidase

**DOI:** 10.64898/2026.05.14.724925

**Authors:** Alexis E. Wilcox, Elizabeth H. Madigan, Catherine J. Andres, Joseph D. Serewicz, Z. Ece Kilic, Tanya S. Freedman, Andrew J. Olive, Caitlyn L. Holmes

## Abstract

*Klebsiella pneumoniae* is a leading cause of pneumonia and bacteremia and is especially dangerous in healthcare settings. Despite massive clinical significance, the mechanisms used by macrophages to kill *K. pneumoniae* are largely unknown. Macrophages are critical for controlling *K. pneumoniae*, as mice lacking monocyte-derived or alveolar macrophages have higher bacterial tissue burdens and mortality. Two prominent mechanisms used by macrophages to kill bacteria are the production of reactive oxygen species (ROS) via the NADPH oxidase NOX2 and reactive nitrogen species (RNS) via the inducible nitric oxide synthase iNOS. Previously, we found that a subset of *K. pneumoniae* factors which enhance bacteremia pathogenesis also support intracellular survival. The degree to which these factors enhance intracellular fitness correlated with resistance against RNS, not ROS. Here, we aimed to define the extent to which macrophage ROS and RNS contribute to *K. pneumoniae* clearance. Using wild-type, *Cybb*^-/-^, and *Nos2*^-/-^ cells, we measured *K. pneumoniae* survival in macrophages lacking such defenses. NOX2 was dispensable for *K. pneumoniae* elimination, and ROS was undetectable in *K. pneumoniae-*infected macrophages. We confirmed that ROS was also undetectable within alveolar-like macrophages, indicating that ROS is dispensable for anti-*K. pneumoniae* defense across macrophage subsets. Instead, iNOS contributed to macrophage clearance of *K. pneumoniae* and likely enhances killing by RNS induction and coordinating a normal cytokine response. Activation of pathways upstream of iNOS may be the most relevant to supporting effective macrophage control of *K. pneumoniae*. This study defines unexpected differential roles for ROS and RNS in macrophage clearance of *K. pneumoniae*.

## Introduction

Sepsis is immune dysregulation that occurs in response to infection and causes immense global morbidity and mortality (1, 2). Bacterial pneumonia and bacteremia are among the most common initiators of sepsis, accounting for >65% of cases (3). The gram-negative species *Klebsiella pneumoniae* is a leading cause of both pneumonia and bacteremia (4–7). *K. pneumoniae* is prominent in healthcare settings, where aging and immunocompromised individuals are at higher risk for infection mortality (8, 9). Furthermore, antibiotic resistance among *K. pneumoniae* strains is increasing, resulting in dangerous infections that are difficult to treat (10). In 2024, the World Health Organization named *K. pneumoniae* as the leading bacterial priority pathogen due to this high virulence and antimicrobial resistance (11). Given the clinical relevance of *K. pneumoniae* and its significant impact on global health, it is critical to understand host-pathogen interactions during these infections.

*K. pneumoniae* infections are highly inflammatory. In murine pneumonia, inflammatory cytokines, such as TNFα, are significantly elevated in the lungs shortly after infection (12). Infected lung tissue also has substantial recruitment of neutrophils and monocyte-derived cells (12, 13). Macrophages are critical for host defense against *K. pneumoniae*, as mice lacking monocyte-derived or alveolar macrophages have higher bacterial burdens in lung tissue and reduced survival during infection (14–16). Since monocyte-derived macrophages are present across infected tissues, understanding how these cells kill *K. pneumoniae* is necessary for defining how this pathogen can be contained across tissues, even after dissemination has occurred. For alveolar macrophages, defining bacterial evasion mechanisms may illuminate opportunities for preventing the dissemination of disease. Despite the importance of macrophages in controlling infection, mechanisms that support effective clearance of internalized *K. pneumoniae* are minimally described.

Macrophages typically internalize pathogens into phagosomes. In this compartment, bacteria are exposed to reactive oxygen species (ROS) and reactive nitrogen species (RNS), two prominent mechanisms used to kill internalized microbes (17, 18). The phagocyte NADPH oxidase (NOX2) assembles onto the phagosome membrane and releases reactive oxygen species (ROS) into the compartment. In parallel, cytosolic inducible nitric oxide synthase (iNOS) generates reactive nitrogen species (RNS), which diffuse across the phagosome membrane to reach bacteria. Although the primary product of NOX2 is superoxide (O_2_ ^-^) and the primary product of iNOS is nitric oxide (NO^-^), these reactive species can form other oxygen and nitrogen-based stressors, such as hydrogen peroxide or peroxynitrite. Such agents interact with various bacterial targets. For example, hydrogen peroxide damages DNA while RNS inhibits bacterial respiration (19). Thus, the distinct targets of ROS and RNS indicate that NOX2 and iNOS may have different contributions to immune stress against *K. pneumoniae,* but this has not been explored in the context of macrophages.

Previously, we used a panel of 53 *K. pneumoniae* mutants with fitness defects during bacteremia and assessed the survival of each after exposure to extracellular ROS and RNS or the macrophage intracellular environment (20). We discovered that a mutant’s ability to resist macrophage-mediated stress was significantly correlated with the ability to resist nitrosative stress. However, no significant correlation was observed between intracellular fitness and oxidative stress resistance. The extent of *K. pneumoniae* resistance to nitrosative stress was also linked to increased bacteremia fitness, but this trend was not observed for oxidative stress resistance. These data indicate that *K. pneumoniae* resistance to nitrosative stress may be particularly relevant during bacteremia and for intracellular survival. However, whether macrophages specifically require ROS and RNS to eliminate *K. pneumoniae* has not been characterized. As such, the role of these stressors in macrophage defense against *K. pneumoniae* remains unclear.

Here, we define the contributions of macrophage NOX2 and iNOS to the killing of intracellular *K. pneumoniae*. We discovered that ROS and NOX2 are dispensable for macrophage elimination of *K. pneumoniae,* and that this species inhibits the initiation of macrophage ROS. Instead, RNS and iNOS significantly contribute to macrophage killing of *K. pneumoniae.* Enhancing pathways that activate RNS production via iNOS may be promising targets for promoting host control of these dangerous infections.

## Results

### iNOS promotes the killing of intracellular *K. pneumoniae*

To characterize macrophage bactericidal mechanisms against *K. pneumoniae,* we first needed to determine the timeframe in which a significant amount of bacterial killing is present. Using gentamicin protection assays, immortalized bone marrow-derived macrophages (iBMDMs) were exposed to the hypervirulent strain KPPR1 for 1 hour (20). Extracellular bacteria were killed with gentamicin, an antibiotic that does not permeate eukaryotic membranes. Intracellular *K. pneumoniae* was either measured immediately (T0) or after two (T2) and four (T4) hours. Macrophages kill a significant proportion of *K. pneumoniae* as early as two hours post infection, with further killing observed after four hours (Figure 1A, Supplemental Figure 1A).

**Figure 1.**
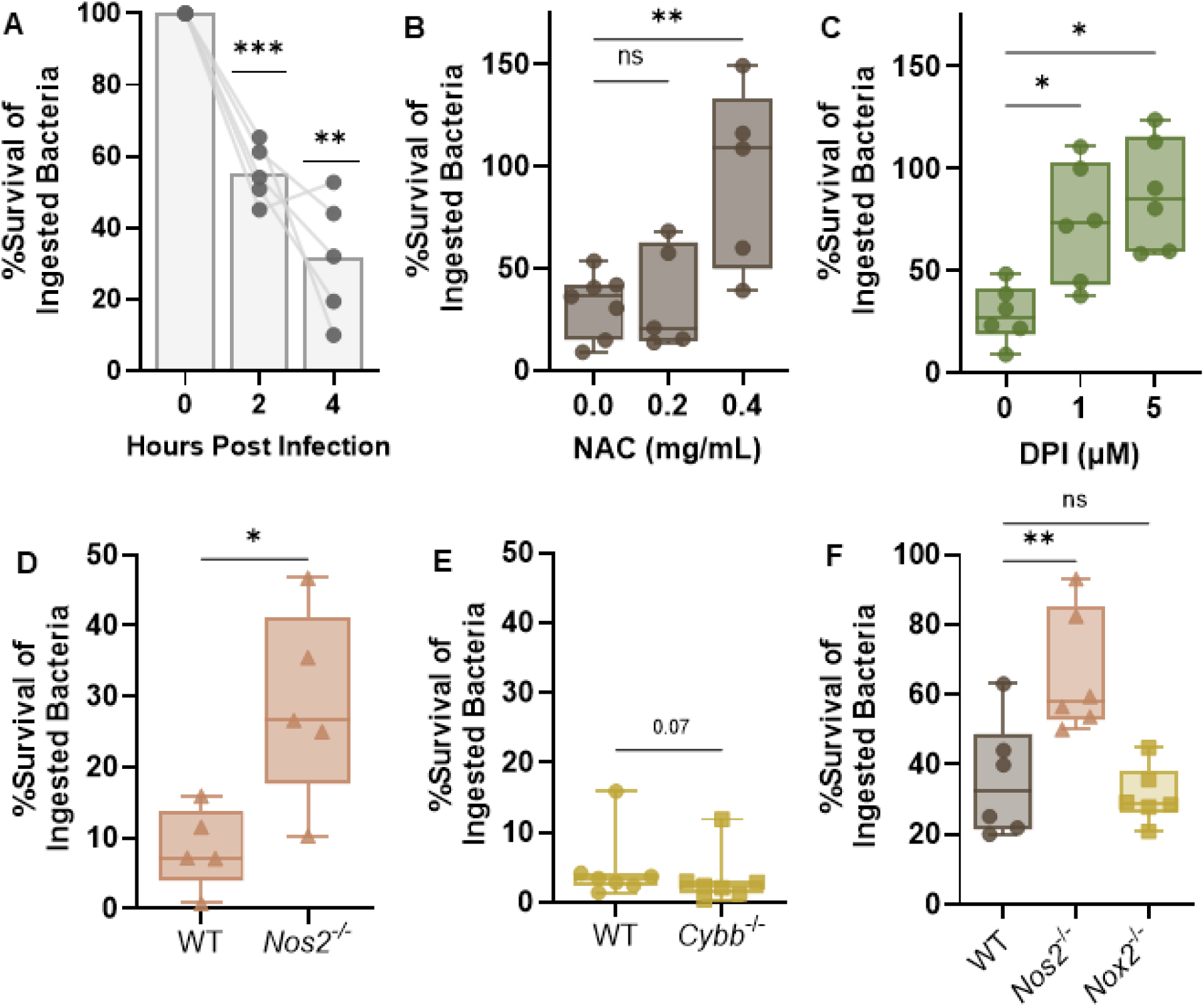
iNOS is required for effective macrophage clearance of *K. pneumoniae*. Macrophages were infected with KPPR1 for one hour prior to the removal of extracellular bacteria with gentamicin treatment. (A) Abundance of intracellular bacteria was assessed immediately (T0), after two hours (T2), or after four hours (T4) and displayed as percent survival, calculated as: (CFU/T0 CFU)*100. \*\*\**p<*0.001, \*\**p*<0.01 by one-way ANOVA. (B) N-acetylcysteine (NAC) or (C) diphenyleneiodonium (DPI) were used to broadly inhibit reactive species or NOX2 and iNOS activity, respectively, at the indicated concentration prior to assessment of KPPR1 intracellular survival. For (B-C), \**p*<0.05*, **p*<0.01 by paired one-way ANOVA. Primary bone marrow-derived macrophages from WT, (D) *Nos2^-/-^*, or (E) *Cybb^-/-^* mice were assessed for the ability to kill intracellular KPPR1. (F) The results for WT, *Nos2^-/-^*, and *Cybb^-/-^*cells were validated using immortalized bone marrow-derived macrophages. For (D-E) \**p*<0.05 by paired *t-*test and *p* values <0.1 are displayed as text; for (F) \*\**p<*0.01 by paired one-way ANOVA comparing each mutant macrophage to WT. For all, n=5-7 trials with symbols representing independent experiments. In (B-F), all values represent the T4 timepoint. For (A), the top of the bar indicates the mean value. For box plots in (B-F), box boundaries represent the 25^th^ and 75^th^ percentile ranges, the middle line represents the median value, and the whiskers represent the minimum and maximum values.

Next, we tested the relative susceptibility of *K. pneumoniae* to killing by distinct reactive species that are generated by the host in the intracellular space. We exposed three strains of *K. pneumoniae* to either hydrogen peroxide, xanthine oxidase and xanthine (generates superoxide and hydrogen peroxide), DETA NONOate (generates nitric oxide), or SIN-1 (generates peroxynitrite) (21, 22). *K. pneumoniae* was susceptible to killing by each type of reactive species. The strain KPPR1 was slightly more sensitive to peroxynitrite than the classical strain Kp4819, but both strains were highly sensitive to killing by this stressor (Supplemental Figure 2).

While *K. pneumoniae* is susceptible to killing by oxidative and nitrosative stress (20), it is unknown whether macrophages specifically leverage ROS and RNS to kill intracellular *K. pneumoniae*. To determine the role of general reactive species in macrophage interactions with *K. pneumoniae*, we broadly neutralized these stressors by adding the antioxidant N-acetylcysteine (NAC) to the media of infected cells (23). At low NAC concentrations (0.2mg/mL), minimal alterations in macrophage killing were observed, but *K. pneumoniae* survival significantly increased at higher NAC concentrations (Figure 1B, Supplemental Figure 1B). NAC had minimal direct effects on *K. pneumoniae* viability as bacterial growth was observed when macrophages were absent (Supplemental Figure 1C). Next, we assessed whether inhibiting sources of ROS and RNS influenced macrophage killing of *K. pneumoniae.* Prior to infection, cells were treated with diphenyleneiodonium (DPI), a chemical inhibitor of flavoenzymes like the NOX2 complex and iNOS (24, 25). *K. pneumoniae* survival was significantly increased after DPI treatment (Figure 1C, Supplemental Figure 1D). While DPI did not exhibit bactericidal properties over a four hour period, high concentrations dampened *K. pneumoniae* replication in media alone (Supplemental Figure 1E). Thus, to clear internalized *K. pneumoniae*, macrophages leverage reactive species that likely originate from flavoenzymes.

To specifically test the involvement of macrophage NOX2 and iNOS in the clearance of *K. pneumoniae*, we generated primary BMDMs from WT mice and from mice lacking the genes *Cybb* and *Nos2* (26). Cybb is a critical component of the NOX2 complex, and in its absence the phagocyte NADPH oxidase cannot assemble; *Nos2* encodes the iNOS enzyme. In the absence of *Nos2*, *K. pneumoniae* survival was significantly elevated compared to WT cells (Figure 1D, Supplemental Figure 3A). In contrast, no difference was observed in *K. pneumoniae* survival between WT cells and those lacking Cybb (NOX2), as both were highly effective at killing *K. pneumoniae* (Figure 1E, Supplemental Figure 3B). We validated these results using iBMDMs sourced from different mice and confirmed that *K. pneumoniae* intracellular survival is significantly increased in cells lacking iNOS (Figure 1F, Supplemental Figure 3C). This trend remains after 24 hours post infection (Supplemental Figure 3C). Notably, we observed minimal differences in *K. pneumoniae* phagocytosis by WT, *Nos2^-/-^,* and *Cybb^-/-^* cells, measured by the abundance of KPPR1 internalized after 1 hour (Supplemental Figure 3D). We also confirmed that WT iBMDMs promote significant induction of *Nos2* in response to KPPR1 and an *E. coli* control (Supplemental Figure 3E). Lastly, macrophage killing of *K. pneumoniae* was not affected by cellular priming prior to infection, as IFNγ-primed BMDMs had similar KPPR1 bactericidal activities compared to untreated cells (Supplemental Figure 3F-G). This is potentially due to robust macrophage inflammation that occurs shortly after *K. pneumoniae* infection.

Collectively, these data confirm that monocyte-derived macrophages leverage iNOS to kill intracellular *K. pneumoniae* and that the NADPH oxidase is dispensable for this process.

### iNOS is required for normal inflammatory signaling in the presence of *K. pneumoniae*

To test whether normal *K. pneumoniae*-induced inflammation is dependent upon iNOS, we collected cellular supernatant from WT and *Nos2^-/-^*primary BMDMs at the beginning and end of the infection. Cytokine abundance in the supernatant was determined by ELISA. Both WT and *Nos2^-/-^* BMDMs had a significant increase in TNFα secretion during *K. pneumoniae* infection (Figure 2A). *Nos2^-/-^* cells trended towards a higher TNFα response after four hours, but this may be due to elevated presence at baseline. *Nos2^-/-^* cells displayed elevated release of pro-inflammatory MCP-1 and IL-6 in response to *K. pneumoniae*, even though these cytokines were minimally detected at baseline (Figure 2B-C). However, we did not detect increased abundance across all inflammatory cytokines during infection. IL-12p40, the active subunit of IL-12 and IL-23, was modestly released in response to *K. pneumoniae* but there was no detectable difference between WT and *Nos2^-/-^*cells (Figure 2D). IL-10 is considered an anti-inflammatory cytokine, but its expression is largely regulated by inflammatory responses. Thus, we were not surprised that *Nos2*^-/-^ macrophages released more IL-10 during *K. pneumoniae* infection than WT cells (Figure 2E). Neither WT nor *Nos2*^-/-^ macrophages elicited IL-1β during these infections as all supernatants were below the limit of detection (data not shown). Collectively, *Nos2*^-/-^ macrophages could be less effective at killing *K. pneumoniae* due to a hyper-inflammatory response that obscures normal killing. The increased inflammation in *Nos2^-/-^*cells may also be due to a higher abundance of internalized *K. pneumoniae* resulting from defects in intracellular killing in the absence of iNOS. These data reveal that iNOS is responsible for normal cellular defense and cytokine release by macrophages in response to *K. pneumoniae*. Thus, iNOS may contribute to *K. pneumoniae* clearance through mechanisms linked to both RNS production and normal immune orchestration.

**Figure 2.**
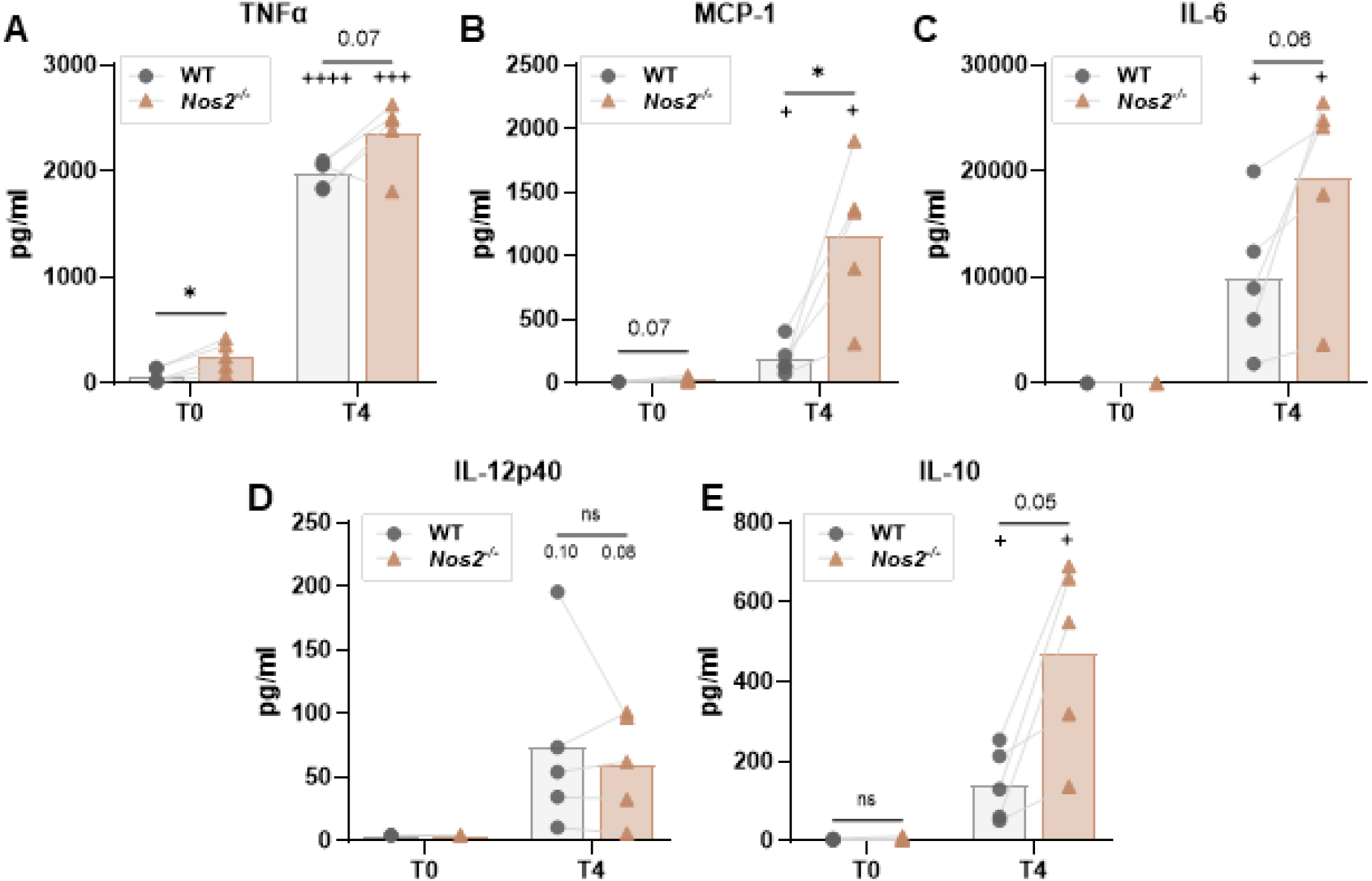
iNOS is required for normal cytokine production in the presence of *K. pneumoniae.* Wild-type and *Nos2*^-/-^ primary bone marrow-derived macrophages were infected with KPPR1 for one hour prior to the removal of extracellular bacteria with gentamicin. Supernatant was collected after gentamicin treatment (T0) or four hours post-infection (T4). Macrophage supernatants were evaluated for cytokine abundance using ELISAs for (A) TNFα, (B) MCP-1, (C) IL-6, (D) IL-12p40, and (E) IL-10. \**p<*0.05 by paired *t-*test; +*p*<0.05, +++*p*<0.001, ++++*p*<0.0001 by paired *t-*test comparing cytokine abundance at T0 vs. T4 for each cell type. For all, *p* values <0.1 are displayed as text; n=5 trials with symbols representing independent experiments. The top of the bar indicates the mean value.

### Macrophage ROS bursts are not detectable in response to intracellular *K. pneumoniae*

We were surprised to observe that NOX2 was dispensable for the intracellular killing of *K. pneumoniae*, given that this is a potent anti-bacterial effector used against many different pathogens (20, 26). Therefore, we measured the extent to which *K. pneumoniae*-infected macrophages generate a ROS burst. *K. pneumoniae* is a remarkably diverse species consisting of two major pathotypes, hypervirulent and classical strains, which can vary in their interactions with the host (13, 27). We assessed the hypervirulent strain KPPR1 and two classical strains: NJST258_2 (ST258), which contains a carbapenem resistance plasmid (28), and Kp4819, which was isolated from the gut of a hospitalized patient (29). First, broad reactive oxygen radicals were measured with the probe 2’,7’-dichlorofluorescin diacetate (DCFDA), a fluorescent dye that tags intracellular redox species. Untreated iBMDMs had minimal ROS at baseline, and ROS were detectable when cells were treated with pyocyanin, a phenazine which stimulates these bursts (30) (Figure 3A). As a separate control, the signal resulting from pyocyanin treatment was confirmed to result from reactive species, as it could be quenched with the antioxidant NAC.

**Figure 3.**
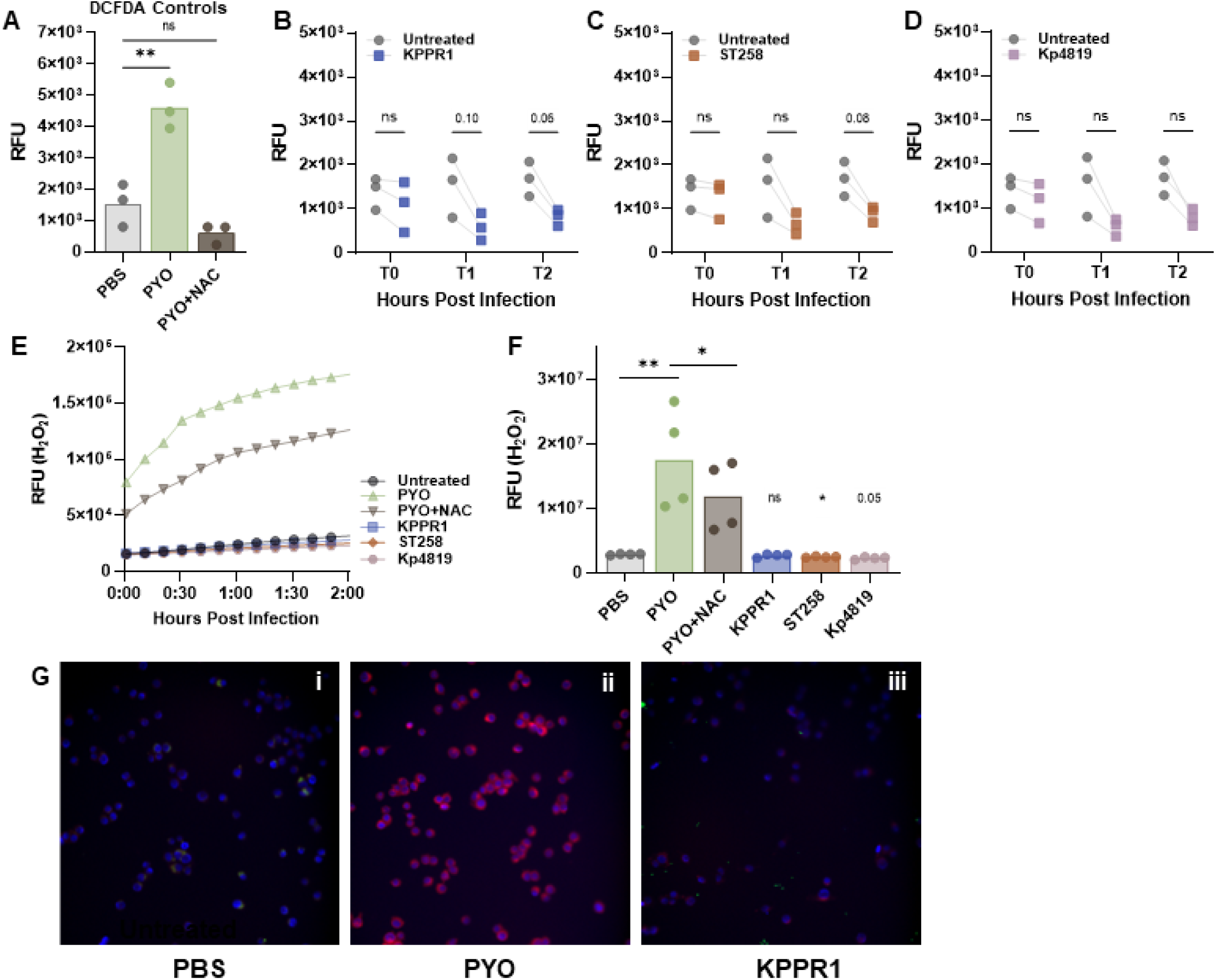
Monocyte-derived macrophages elicit no detectable ROS burst in response to *K. pneumoniae*. (A) Immortalized bone marrow derived macrophages were untreated (PBS), exposed to pyocyanin, or treated with pyocyanin+N-acetylcysteine (NAC). Relative fluorescence (RFU) of controls was measured 1 hour post infection. Macrophages were infected with (B) KPPR1, (C) ST258, or (D) Kp4819 and stained with DCFDA. (E) Macrophages were treated as above for Amplex Red staining. RFU was measured every 10 minutes for two hours, and the average RFU for four trials is displayed. (F) Area under the curve (AUC) was measured for the data in (E). (G) Macrophages were (i) untreated, (ii) treated with pyocyanin, or (iii) infected with KPPR1-chromoGFP then stained with CellROX (red) and NucBlue (blue). Representative images are displayed for one of three trials. For (A) \*\**p*<0.01 by one-way ANOVA; for (B-D), no comparisons were statistically signifi ant by paired *t-*test and lines connect data from the same trial. For (F), comparisons with lines are paired *t-* tests comparing the control groups and *\*p<*0.05, \*\**p*<0.01; the AUC for cells infected with KPPR1, ST258, and Kp4819 were compared to untreated cells using a paired one-way ANOVA and *\*p*<0.05. For all panels, comparisons with *p*<0.1 are indicated in text and n=4 independent trials. In (A) and (F), the top of the bar indicates the mean value; in all graphs, symbols represent independent trials.

Macrophage ROS was measured via DCFDA after BMDMs were infected with KPPR1 (Figure 3B), ST258 (Figure 3C), and Kp4819 (Figure 3D). ROS signal was measured shortly after infection (T0), one (T1), or two hours post-infection (T2) since macrophage anti-bacterial effectors are present at this time. Macrophage ROS was undetectable in response to each *K. pneumoniae* strain, even though classical strains were significantly more internalized than the hypervirulent strain (Supplemental Figure 4A). Instead, *K. pneumoniae* infected macrophages exhibited trends towards lower ROS than untreated cells, although this did not reach statistical significance. We then assessed whether this phenotype was specific to *K. pneumoniae* and measured ROS in macrophages infected with two other Gram-negative species, *Escherichia coli* or *Pseudomonas aeruginosa*. Macrophages generated substantial ROS bursts in response to infections with these species (Supplemental Figure 4B-C), further emphasizing a unique ability of *K. pneumoniae* to dampen host ROS. Priming macrophages with IFNγ increased baseline ROS, but stimulated cells did not produce a ROS burst upon *K. pneumoniae* infection, again confirming the ROS blocking phenotype (Supplemental Figure 4D-E). To broadly define *K. pneumoniae* factors that contribute to ROS blocking, we treated macrophages with UV-killed KPPR1 and again observed minimal ROS signal. Thus, the ability of *K. pneumoniae* to block ROS is likely due to a passive feature of the bacteria.

Next, we confirmed the DCFDA results using a secondary probe, Amplex Red, which measures the accumulation of extracellular hydrogen peroxide (31). ROS detection via Amplex Red was feasible as macrophages stimulated with pyocyanin elicited a significantly higher signal than untreated cells, and the signal was quenched with NAC (Figure 3E-F). As with DCFDA, macrophages did not elicit detectable hydrogen peroxide release over a two hour period in response to any *K. pneumoniae* strain. Again, *K. pneumoniae* strains had seemingly lower ROS signal than untreated cells, and this reached statistical significance for the strain ST258 when measured with Amplex Red (Figure 3F).

We next visualized whether individual macrophages elicit ROS that would be undetectable within a population-based plate reader assay like DCFDA or Amplex Red. Using the probe CellROX, which generates a fluorescent signal when in contact with hydroxyl species, we visualized ROS within KPPR1-infected cells. Untreated macrophages elicited no detectable ROS signal, but ROS could be visualized after pyocyanin treatment (Figure 3Gi-ii). Macrophages infected with KPPR1 demonstrated no detectable ROS signal on an individual cellular level (Figure 3Giii). These collective data using three independent techniques reveal that macrophages do not elicit a detectable ROS burst in response to *K. pneumoniae*. These results are supported by the finding that NOX2 activity is dispensable for the eradication of intracellular *K. pneumoniae*.

### *K. pneumoniae* does not inactivate NOX2 signaling

Since infected macrophages did not elicit a detectable ROS burst, we tested whether this was due to *K. pneumoniae* inhibition of the active, NOX2 signaling cascade. First, macrophages were infected with *K. pneumoniae* for one hour to allow time for host-pathogen interactions to initiate. Then, NOX2 activity was induced using the protein-kinase-C (PKC) isotype activator phorbol 12-myristate 13-acetate (PMA). PKC activates *p*47*^phox^* (NCF1), causing rapid assembly and induction of NOX2 upon stimulation (32, 33). We reasoned that if *K. pneumoniae* inhibited NOX2 activity, then infected macrophages would have minimal ROS signal after PMA treatment. Uninfected macrophages displayed elevated ROS after PMA treatment, and NAC quenched the signal (Figure 4A-B). As before, no ROS signal was detected in cells treated with *K. pneumoniae* alone (for KPPR1, ST258, or KP4819). However, infected macrophages which were treated with PMA did indeed produce a substantial ROS burst. Although *K. pneumoniae* elicited minimal ROS production, iBMDM *Cybb* transcript levels were comparable during infection with *K. pneumoniae* and *E. coli* infected cells, and Cybb protein was detectable in BMDMs throughout infection (Supplemental Figure 4F-H). As expected, Cybb was not detectable in *Cybb^-/-^*iBMDMs. Collectively, these data indicate that NOX2 assembly and activity can still occur in the presence of *K. pneumoniae* given a sufficient stimulus. Therefore, *K. pneumoniae* may prevent the initial assembly of the NOX2 complex.

**Figure 4.**
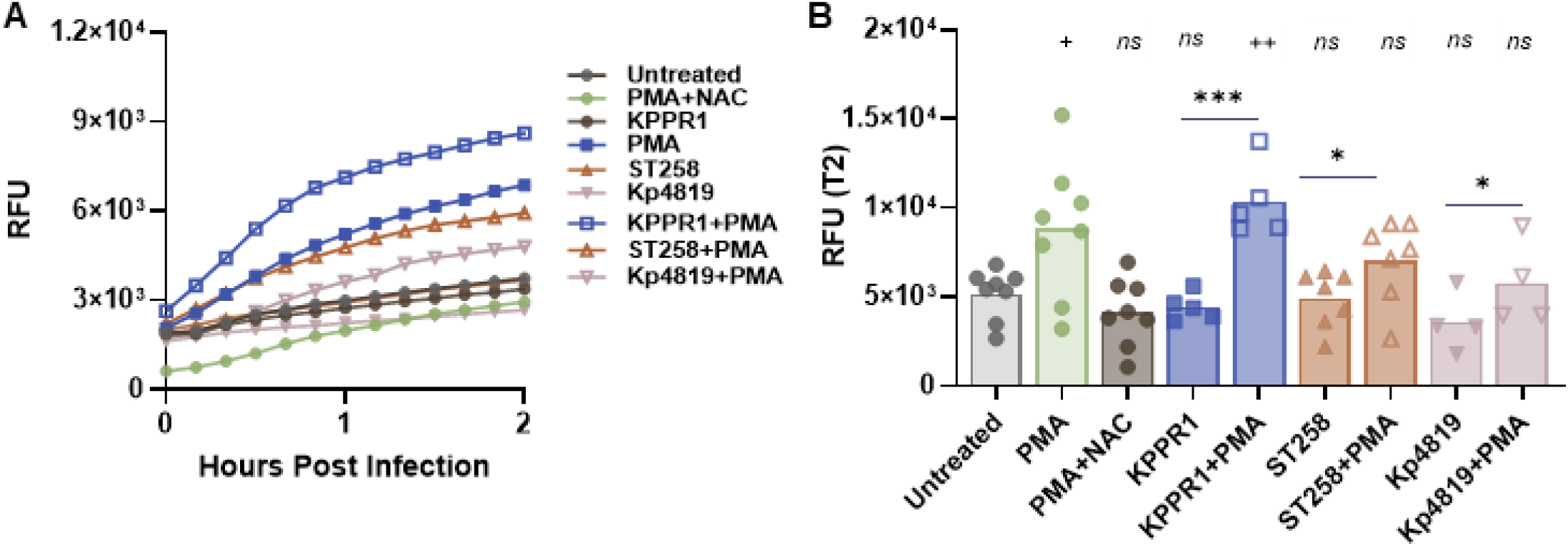
*K. pneumoniae* does not block induced NADPH signaling. Primary bone marrow derived macrophages were infected with KPPR1, ST258, or Kp4819 and then either left untreated or stimulated with PMA. (A) The DCFDA relative fluorescence (RFU) of each well, including controls (untreated, PMA, and PMA+NAC), was measured every ten minutes for two hours. The mean value is displayed for each group. (B) The endpoint relative fluorescence units (RFU) at T2 is displayed for the curves in (A). For (B), +*p*<0.05 and ++*p*<0.01 by one-way ANOVA comparing each group to the untreated control; *\*p<*0.05 and \*\*\**p<*0.001 by paired *t-*test comparing the signal from macrophages infected with each *K. pneumoniae* strain with and without PMA stimulus. The top of the bar indicates the mean value; n=4-7 and symbols represent independent trials.

### *K. pneumoniae* evades ROS generation in alveolar-like macrophages

Since ROS generation was undetectable in *K. pneumoniae*-infected monocyte-derived cells, we tested whether this was consistent across a second, distinct macrophage lineage. Both monocyte-derived and alveolar macrophages are important for *K. pneumoniae* infection control, yet have fundamentally different host-pathogen interactions, as they arise from separate stem-cell progenitors and have distinct baseline functions (14–16, 34, 35). To investigate whether *K. pneumoniae-*infected alveolar-like macrophages lack an ROS burst, we leveraged a model of fetal liver-derived alveolar-like macrophages (FLAMs), (34). FLAMs were less phagocytic of KPPR1 than BMDMs but had significant bactericidal activity after two hours (Figure 5A-B).

**Figure 5.**
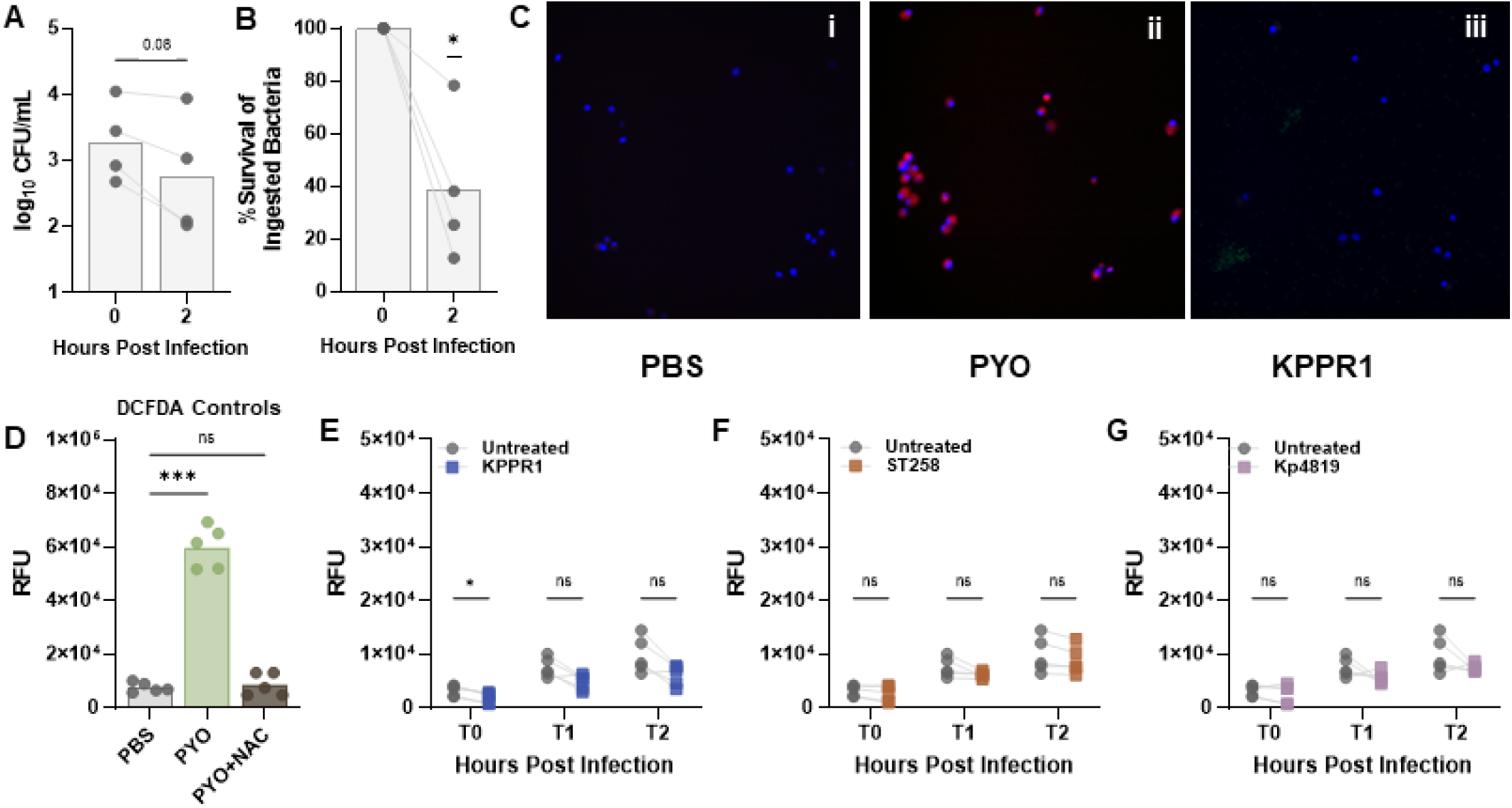
Alveolar-like macrophages elicit no detectable ROS burst in response to *K. pneumoniae*. Fetal liver-derived alveolar-like macrophages (FLAMs) were infected with KPPR1 for one hour prior to the removal of extracellular bacteria with gentamicin treatment. The abundance of intracellular bacteria was assessed immediately (T0) or after two hours (T2). In (A), the abundance of intracellular KPPR1 is displayed as log_10_ CFU/mL with significance assessed by paired *t-*test and the *p* value indicated by text. In (B), the results from (A) are displayed as percent survival, calculated as: (T2 CFU/T0 CFU)*100. \**p<*0.05 by one sample *t-*test with a hypothetical value of 100. (C) FLAMs were (i) untreated, (ii) treated with pyocyanin, or (iii) infected with KPPR1-chromoGFP (green) then stained with CellROX (red) and NucBlue (blue). Representative images are displayed for one of three independent trials. (D) FLAMs were left untreated, exposed to pyocyanin, or treated with pyocyanin+N-acetylcysteine. Relative fluorescence (RFU) of controls was measured 1 hour post infection. \*\*\**p*<0.001 by one way ANOVA. FLAMs were infected with (E) KPPR1, (F) ST258, or (G) Kp4819 and stained with DCFDA. \**p*<0.05 by paired *t-*test and lines connect values from the same trial. For all, symbols represent independent trials; in (A), (B), and (D), the tops of bars indicate mean values.

FLAMs were infected with KPPR1, and ROS was visualized with CellROX. Untreated FLAMs produced minimal ROS in comparison to pyocyanin-treated FLAMs (Figure 5Ci-ii). As with BMDMs, KPPR1-infected FLAMs had no detectable ROS burst via microscopy (Figure 5Ciii). These data were confirmed with DCFDA; FLAMs generated a robust ROS burst following pyocyanin treatment that could be quenched with NAC (Figure 5D). Again, no ROS signal was detected across KPPR1 (Figure 5E), ST258 (Figure 5F), or Kp4819 (Figure 5G), even though FLAMs phagocytized substantially more ST258 and Kp4819 than KPPR1 (Supplemental Figure 5A). As with monocyte-derived macrophages, priming of FLAMs with IFNγ prior to *K. pneumoniae* infection did not alter bactericidal activity (Supplemental Figure 5B-C). Although priming with IFNγ upregulated ROS production in uninfected cells, this activation did not result in ROS production following *K. pneumoniae* infection (Supplemental Figure 5D-E). As with BMDMs, FLAMs had significant *Nos2* and *Cybb* transcription following both *K. pneumoniae* and *E. coli* infection (Supplemental Figure 5F-G). FLAMs also had detectable Cybb protein throughout infection with both species (Supplemental Figure 5H-I). Notably, FLAM Cybb levels increased over time during *Kp* infection, but remained steady in iBMDMs, further highlighting differences in host-pathogen interactions in different macrophage subsets (Supplemental Figure 4G-H, 5H-I). These data confirm that two distinct macrophage lineages do not elicit detectable ROS upon infection with diverse strains of *K. pneumoniae*.

## Discussion

Here, we tested the contributions of two macrophage antibacterial effector mechanisms, oxidative and nitrosative stress, to the killing of intracellular *K. pneumoniae*. Our data reveal that macrophages utilize iNOS, rather than the phagocyte NADPH oxidase NOX2, as a *K. pneumoniae* clearance mechanism. iNOS likely supports *K. pneumoniae* elimination both through RNS production and by orchestrating a normal immune response. Macrophages infected with *K. pneumoniae* demonstrated no detectable ROS burst, further supporting the dispensability of NOX2 in these interactions. *K. pneumoniae* likely blocks NOX2 assembly, rather than interfering with signaling once it occurs. Importantly, these findings are applicable across macrophage lineages and *K. pneumoniae* strains, highlighting novel host-pathogen interactions that are relevant in multiple infection types.

It is well-established that NOX2 is critical for host defense against *K. pneumoniae*. Our research group and others have reported that mice lacking functional NOX2 have higher bacterial burdens in the lung and lower survival following *K. pneumoniae* infection (36–38). Mice lacking NOX2 also have altered *K. pneumoniae* dissemination patterns from the lung to the blood, seemingly due to ROS-dependent control of the initial KPPR1 founding population (37). However, ROS is not equally effective in anti-*K. pneumoniae* defense across immune subsets. A previous study found that the NOX2 inducing protein SKAP2 is required in neutrophils, not macrophages, to support KPPR1 clearance *in vivo* (38). In accordance with this study, our data show that macrophages do not require ROS to kill intracellular *K. pneumoniae.* Thus, neutrophils are likely the prominent source of oxidative stress during *K. pneumoniae* infections.

Instead, our data reveal that iNOS is required for effective macrophage control of *K. pneumoniae*. On average, BMDMs derived from *Nos2^-/-^* mice had ∼5 fold lower KPPR1 killing than WT cells. iNOS is also required for *in vivo* defense against KPPR1, as mice treated with the iNOS inhibitor L-NAME had significantly lower survival and elevated bacterial burden in the lung and blood (39). Alveolar macrophages treated with L-NAME show reduced *K. pneumoniae* killing, demonstrating that this mechanism is used across multiple macrophage lineages (39). Additionally, human alveolar macrophage clearance of *K. pneumoniae* is substantially enhanced upon treatment with surfactant protein A, which elevates cellular nitric oxide (21). Although *Nos2^-/-^*cells were significantly less bactericidal than WT cells, they still cleared ∼1.5 logs of internalized KPPR1 at a 24-hour timepoint. These results indicate that iNOS is required for effective early defense against intracellular *K. pneumoniae* but that later defenses also potentiate killing. We also identified roles for iNOS in defense against *K. pneumoniae* beyond direct bactericidal activity as cytokine production was altered in the absence of iNOS. Our data aligns with previous reports that iNOS partially dampens M1 polarization, such that cells lacking iNOS may be more likely to transition to an inflammatory state (40).

*K. pneumoniae* bacteremia fitness factors have multiple overlapping roles with the ability to resist oxidative, nitrosative, and macrophage-mediated stress (20). The extent to which bacteremia fitness mutants resist RNS is significantly correlated with their ability to resist macrophage-mediated stress, yet there is no substantial link between resistance against oxidative stress and resistance to the intracellular environment. Our results in the present study support these findings. The contributions of iNOS, and the dispensability of NOX2 further define that macrophages leverage RNS to elicit stress on intracellular *K. pneumoniae*. Nitric oxide can have significant impacts on *K. pneumoniae,* including the alteration of capsule and biofilm formation (41). *K. pneumoniae* also generates inherent RNS in response to external oxidative and nitrosative stress (20), meaning that these stressors could potentiate bacterial death by eliciting self-production of this molecule. Accordingly, *K. pneumoniae* employs multiple mechanisms to resist nitrosative stress. Various *K. pneumoniae* regulators and metabolic genes enhance resistance against RNS, although studies identifying these RNS resistance factors have tested a limited number of factors (20, 42). Further work should identify the broader scope of *K. pneumoniae* genes regulating resistance against nitrosative stress as targeting these pathways could support more effective bacterial clearance by macrophages.

In support of the finding that NOX2 is dispensable for macrophage clearance of *K. pneumoniae*, we identified no detectable ROS burst within infected cells. Our monocyte-derived and alveolar-like macrophages displayed healthy morphology and bactericidal capabilities. We were also able to detect ROS within both cell types through positive controls, confirming that the cells were capable of producing ROS bursts. However, infected cells from neither lineage elicited a detectable ROS signature. We further confirmed these phenotypes in IFNγ-primed cells, which had increased baseline ROS but continued to display no induction of oxidative bursts or enhanced bactericidal activity during *K. pneumoniae* infection. Because *K. pneumoniae* infected cells become highly inflammatory shortly after infection, we believe that pre-treatment or polarization to the M1 state is redundant for the methodology used in this study. Since *K. pneumoniae* did not dampen ROS when initiated via PMA treatment, we believe that initial NOX2 assembly within macrophages is prevented rather than *K. pneumoniae* playing an active role in inhibiting ROS signaling. Of note, both live and dead KPPR1 were capable of blocking ROS bursts in comparison to known ROS stimuli such as *E. coli* and *P. aeruginosa*. These results indicate that ROS blocking by *K. pneumoniae* is likely a passive feature, and does not require bacterial viability. This phenotype could be due to complex cell surface features, and the identity of such factors are the focus of ongoing studies.

The absence of ROS within macrophages infected with diverse isolates indicates that *K. pneumoniae* has conserved mechanisms for thwarting this host defense. The *K. pneumoniae* species contains two major pathotypes containing hypervirulent and classical strains. Hypervirulent strains are characterized by enhanced capsule production, siderophores, and invasive disease in healthy patients (43). Classical strains are associated with multidrug resistance and outbreaks in healthcare settings. These differences can alter interactions with the host including bacterial recognition, phagocytosis, and induced inflammation (13, 44). To account for these differences, our study incorporated three diverse isolates, one hypervirulent (KPPR1) and two classical (ST258, Kp4819) strains. As no strain of *K. pneumoniae* in this study elicited macrophage ROS across two distinct lineages, our results suggest a widely applicable phenotype of ROS blocking by this pathogen.

We recognize that this study has limitations including the use of *in vitro* assays and the investigation of one immune subset. Within infected tissue, *K. pneumoniae* is exposed to a cytokine-rich environment and other cell-cell interactions that are not accounted for with our *in vitro* approach (45, 46). However, we believe the current approach is sufficient to detect distinct contributions from NOX2 and iNOS in this process. Our work focused on murine macrophages, but it is possible that iNOS has differential contributions within human cells. While our study was focused on the role of NOX2 and iNOS in macrophage killing of *K. pneumoniae*, it did not extend these characterizations to other innate immune subsets such as neutrophils. NOX2 is an established source of anti-*K. pneumoniae* defense by neutrophils, but the extent to which neutrophil iNOS supports *K. pneumoniae* killing will be the focus of future work.

Collectively, our study provides new insight into the mechanisms used by macrophages to eliminate intracellular *K. pneumoniae*. We discovered surprising, divergent roles for NOX2 and iNOS in macrophage defense against *K. pneumoniae* with iNOS being required for effective killing of intracellular *K. pneumoniae*. Thus, pathways upstream of iNOS that elicit this response may be highly relevant for enhancing host defense against this dangerous pathogen.

## Materials and Methods

### Bacterial Strains and Materials

All materials and reagents were sourced from ThermoFisher (Fisher Scientific, Waltham, MA) unless stated otherwise. The *K. pneumoniae* strains used in this study are: ATCC 43816 (KPPR1), 30684/NJST258_2 (ST258), and Kp4819 (28, 47, 48). The fluorescent strain KPPR1-chromoGFP expresses constitutive GFP expression from the chromosome and was constructed previously (20).

*K. pneumoniae* strains were stored at −80°C and maintained on Lysogeny broth (LB, BP1425500) plates for up to 3 weeks at 4°C. Prior to experiments, bacteria were cultured by inoculating 2 mL of LB broth (BP1426500) with a single colony and grown overnight, shaking at 37°C. *K. pneumoniae* overnight cultures were washed with phosphate-buffered saline (PBS, MT21040CV) and adjusted to the appropriate experimental concentration based on OD_600_ density. To kill bacteria prior to experimentation, the inoculum was incubated under UV light for 1 hour. Quantitative culture was used to verify the bacterial input for all experiments.

### Cell Culture and Primary Cell Isolation

Immortalized bone marrow derived macrophages (iBMDMs) were initially derived from C57BL/6 mice (49, 50). iBMDMs were cultured using DMEM (MT10017CV)+10% heat-inactivated fetal bovine serum (FBS; MT35011CV)+1% penicillin-streptomycin (MT30009CI) on 100mm tissue culture treated dishes (FB012924) until 60-90% confluency. For passaging, cells were rinsed with PBS and lifted from the dish using with 2mM ethylenediaminetetraacetic acid (EDTA; BP2482100). Cells were washed, resuspended in DMEM+10% FBS, and quantified using a hemacytometer with viability assessed by trypan blue (15-250-061) exclusion. All cells used in this study demonstrated >90% viability. Once quantified, cells were either seeded for experimentation or passaged for continued proliferation. iBMDMs used for experiments were between 5-15 passages.

Primary BMDMs (pBMDMs) were isolated from the femurs of mice 6-18 weeks. Bone marrow was flushed with DMEM+10% heat inactivated FBS+1% penicillin-streptomycin and passed through a 40 micron cell strainer (08-771-1). Cells were washed and resuspended in DMEM+10% heat inactivated FBS+1% penicillin-streptomycin+15% L-cell supernatant. Nucleated cells were counted on a hemacytometer and seeded into non-tissue culture treated 100 mm dishes at a density of 5 million cells (FB0875712). 3-4 days after initial seeding the media was exchanged. One week after seeding and subsequent proliferation, pBMDMs were harvested by lifting with 2mM EDTA. The cell suspensions were pelleted and resuspended in fresh media prior to quantification. Frozen pBMDM stocks were prepared by aliquoting cells into cryo-grade vials with 10% dimethyl sulfoxide (DMSO; 25-950-CQC). Stocks were placed into foam containers to allow for gradual cooling and moved 24 hours later to liquid nitrogen for long term storage. Once thawed from liquid nitrogen, pBMDMs were seeded for experimentation.

Fetal liver-derived cells were isolated as previously described by plating cells retrieved from the livers from mice at roughly E18 (34, 51). Alveolar-like macrophages (FLAMs) were then generated as previously detailed. FLAMs were cultured in 100mm tissue culture dishes in RPMI (MT10040CV)+10% heat inactivated FBS+1% penicillin streptomycin+30ng/ml recombinant GM-CSF+15ng/ml recombinant TGF (PeproTech, Cranbury, NJ: 3150320UG, 100-21-10UG). FLAMs were seeded at densities no less than 750,000 and no more than 2 million cells per dish and passaged at 60-75% confluency. Cells were lifted from the dishes using 10mM EDTA, washed, and enumerated on a hemacytometer. FLAMs used for experiments were between 5-12 passages.

For all assays in which cells were primed to an inflammatory state prior to experiments, cells were incubated with media containing 200ng/mL IFNγ (StemCell Technologies). Macrophages were kept under the influence of IFNγ from the time of seeding until the experiment concluded.

### Gentamicin Protection Assays

To assess *K. pneumoniae* uptake and intracellular survival within macrophages, gentamicin protection assays were performed as previously described (20). Briefly, iBMDMs were seeded and infected with *K. pneumoniae* at a target ratio of 10 bacteria per cell (MOI 10) in 24 well tissue culture treated dishes (09-761-146). The infection was synchronized by centrifugation at 500xg for five minutes at 4°C. The plate was then incubated at 37°C with 5% CO_2_ for one hour to allow for bacterial uptake. Extracellular bacteria was killed with gentamicin (15750060) at a concentration of 100ug/mL in the appropriate cell culture media for 30 minutes. Afterwards, a subset of wells were washed with 1mL PBS and lysed with 500uL of filter-sterilized 1x saponin (AAJ63209AK), and bacterial uptake assessed by quantitative culture (T0). A separate set of wells were incubated for an additional 2, 4, or 24 hours at 37°C, 5% CO_2_ prior to washing, lysis, and enumeration (T2, T4, T24). Intracellular survival was assessed as the abundance of *K. pneumoniae* surviving at each timepoint: percent survival of ingested bacteria = (CFU T2, T4, or T24/CFU T0)*100. Experiments using chemical inhibitors had the following modifications. N-acetylcysteine (NAC, 20261; Cayman Chemical Company, Ann Arbor, MI) was diluted into cell culture media and added to macrophages at the time of infection at 0.2 or 0.4mg/mL. For diphenyleneiodonium chloride (DPI, 50-176-2440; Sigma-Aldrich, St. Louis, MO), 1 or 5µM/mL was added to macrophages at the time of initial cell seeding, roughly 16 hours prior to the start of the experiment. For all experiments using chemical inhibitors, the ability of *K. pneumoniae* to replicate in the presence of the inhibitor was assessed. *K. pneumoniae* (1×10^7^ CFU) was added to media containing individual inhibitors at the concentrations listed above. Bacterial survival was quantified by sampling wells for enumeration at T0 and T4. For gentamicin protection assays using FLAMs, macrophages were seeded in tissue culture treated 12 well dishes using RPMI+10%FBS. For experiments using primary BMDMs, cells were seeded in DMEM+10% FBS in non-tissue culture treated 6 well dishes.

### ELISA

The supernatants of infected pBMDMs were analyzed for cytokine abundance using ELISA. Macrophages were infected as detailed for gentamicin protection assays. At the T0 and T4 timepoint, 1mL of media was removed and immediately banked at −80°C. Supernatants remained frozen until analysis by the University of Minnesota Cytokine Reference Laboratory. All ELISAs were performed with standard curves and positive and negative controls.

### Measurement of ROS

All ROS detection assays included controls to assess baseline ROS signal (PBS, untreated cells) and baseline *K. pneumoniae* fluorescence (bacteria only). Assays also included positive controls to ensure ROS signal was detectable, either 1mM pyocyanin (601522; Cayman Chemical Company) or 1.6mM phorbol 12-myristate 13-acetate (PMA, 50-288-7125; Cayman Chemical Company) depending on the assay.

To measure intracellular radicals, 2’,7’-dichlorofluorescin diacetate (DCFDA) was used (601520, Cayman Chemical). Macrophages were seeded in a black-walled tissue culture treated 96-well dish (353219) at 5×10^4^ cells/well in the appropriate cell culture media and infected with *K. pneumoniae* to achieve MOI 50. After infection, cells were centrifuged at 500xg for 5 minutes to synchronize infection and incubated at 37°C, 5% CO_2_ for 1 hour. DCFDA staining occurred following the manufacturer’s protocol with the exception that 100µg/mL of gentamicin was added to ROS Staining Buffer to kill extracellular bacteria. After the 30-minute staining incubation, fluorescence was measured using a Synergy H1 Hybrid plate reader with an excitation wavelength of 490nm and an emission wavelength of 530nm. Relative fluorescence units (RFU) were calculated by averaging two technical replicates of infected cells then subtracting background signal generated by bacteria using fluorescence from wells containing bacteria only. For all *K. pneumoniae* strains, this signal was minimal to undetectable.

For Amplex Red assays, BMDMs were seeded in black-walled tissue culture treated 96-well dishes at 5×10^4^ cells per well. Macrophages were infected with *K. pneumoniae* with a target MOI of 50. After infection, cells were centrifuged at 500xg for 5 minutes to synchronize infection and incubated at 37°C, 5% CO_2_ for 1 hour. Cells were washed and Amplex Red stock solution was applied (10mM Amplex Red Reagent, A22188, 10U/mL HRP stock solution, 1X Reaction Buffer) to each well. Technical replicates were run in duplicate and values were averaged. Fluorescence was measured every 10 minutes for 2 hours, with emission detection at 590nm and an absorbance of 560nm using a Synergy H1 Hybrid plate reader.

For CellROX imaging, macrophages were seeded in black-walled tissue culture treated 96-well dishes at 1×10^4^ cells per well in the appropriate cell culture media. Cells were either uninfected, treated with 1mM pyocyanin or infected at an MOI 50 with KPPR1-chromoGFP. Plates were centrifuged at 500xg for 5 minutes to facilitate infection and incubated at 37°C, 5% CO_2._ After incubation, media was removed, and cells were treated with 100µg/mL gentamicin for 30 minutes. Cells were then washed, stained with CellROX Red (C10422) and NucBlue (PI62249) for 30 minutes, and fixed with 4% paraformaldehyde (15714). Cells were imaged within an hour of staining utilizing 20x magnification on a Keyence BZ-X810 fluorescence microscope. Technical replicates were run in duplicate to ensure similar patterns in stained could be observed across wells.

### Immunoblotting and Quantification

Immunoblotting and quantification were performed as described previously (52). Briefly, iBMDMs and FLAMs were infected as described above for gentamicin protection assays. Cell lysates were prepared at T0, T4, and T24 by placing cells on ice and incubating in sodium dodecyl sulfate (SDS) buffer before storage at −20°C. Lysates were separated on a 7% NuPage Tris-Acetate gel (Invitrogen) and transferred to an Immobilon-FL PVDF membrane (EMD Millipore). Revert Total Protein Stain (LI-COR Biosciences) was used to quantify the total protein content in each lane. After destaining, membranes were cut into appropriate sections and blocked 1 hour with Intercept Tris-buffered saline (TBS) Blocking Buffer (LI-COR). After blocking, membranes were incubated overnight at 4°C with Cybb primary antibody (BD Biosciences, 611414) followed by a fluorophore-conjugated secondary antibody (IRDye 800CW donkey anti-mouse IgG, LI-COR Biosciences, 926-32212). Blots were visualized with an Odyssey CLx near-infrared imager (LI-COR), and protein signals were analyzed using ImageStudio Software (LI-COR). The protein signals were background-subtracted, corrected for whole-lane protein content, and shown relative to T0 for each cell type infected with KPPR1 or DH5α. After analysis, images were rotated, cropped, and adjusted in Adobe Photoshop. Corrections for brightness and contrast were applied uniformly to each image, where applicable.

### RNAseq

Cells were infected in the same manner as the gentamicin protection assays, described above. At 0 and 4 hours post infection, cells were washed with PBS and lifted using 2mM EDTA. The cell suspension was pelleted at 500xg for 5 minutes at 4°C and resuspended in 150µL of DNA/RNA Shield (Zymo Research Corporation). Samples were immediately stored at 4°C until processed and analyzed by Plasmidsaurus, according to the Differential Gene Expression Analysis work flow service. All samples were classified by Plasmidsaurus as having high RNA concentration and quality, and each sample contained ∼20 million reads.

### In vitro Chemical Stress Assays

For all in vitro stress assays, 1×10^6^ CFU of KPPR1, Kp4819, and NJST258-2 were incubated at 37°C and 5% CO_2_ with reactive species. Bacteria were incubated with 1mM hydrogen peroxide or 2.5 mM DETA NONOate (82120, Cayman Chemical) for 2 hours as previously described (20). Bacteria were incubated with 1µM 3-Morpholinosydnonimine hydrochloride (SIN-1 hydrochloride) (AB141525, Abcam) or 100nU/mL xanthine oxidase (X2252, Sigma Aldrich) and 10nM xanthine (X7375, Sigma Aldrich) for 30 minutes (21, 22). Bacteria were enumerated by serial dilutions and spot plating on LB agar at the beginning and end of the experiment. Plates were incubated at 27°C and then counted, percent survival was calculated by dividing the T2 by the T0 timepoint.

### Statistical Analysis

Statistical significance was calculated using GraphPad Prism software. Ordinary one-way ANOVA with Dunnett’s multiple comparisons was used to define differences in means among multiple groups in reference to a single mean, one-sample *t*-tests calculated the difference in a group in reference to a theoretical mean, paired or unpaired *t-*tests compared differences among two groups. A *p-*value <0.05 was determined to be statistically significant. All assays were run in at least three independent trials.

## Acknowledgements

Research reported in this publication was supported by the National Institute of Allergy and Infectious Diseases of the National Institutes of Health under Award Number R00AI175481 (CLH), the National Institute for General Medical Sciences of the National Institutes of Health under Award Number R35GM146795 (AJO), and the National Institute of Arthritis and Musculoskeletal and Skin Diseases under Award Number R01AR084525 (TSF). The content is solely the responsibility of the authors and does not necessarily represent the official views of the National Institutes of Health. The authors thank Dr. Mary O’Riordan, Dr. Michael Bachman, Dr. JD Sauer, and Dr. Matthew Freeman for sharing the *Nos2^-/-^* and *Cybb^-/-^* mutants from primary and immortalized BMDM lineages. Aiden Backes, Telmen Gantulga, and Ali Ahmed assisted with phagocytosis assays and *in vitro* assays under the mentorship of AEW, EHM, and JDS. We thank Dr. Julia L. E. Willett for sharing the DH5α and PA14 strains. Jessica Herndon assisted with figure design. Dr. Jay Vornhagen and Dr. Karthik Hullahalli assisted with manuscript feedback.

## Author Contributions

AEW: data curation, formal analysis, investigation, methodology, validation, visualization, writing – original draft, writing – reviewing and editing

EHM: formal analysis, investigation, methodology, validation, writing – original draft, writing – reviewing and editing

CJA: formal analysis, investigation, methodology, writing – original draft, writing – reviewing and editing

JDS: formal analysis, investigation, writing – original draft, writing – reviewing and editing

ZEK: formal analysis, investigation, methodology, writing – reviewing and editing

TSF: formal analysis, funding acquisition, methodology, supervision, validation, writing – reviewing and editing

AJO: funding acquisition, methodology, validation, writing – original draft, writing – reviewing and editing

CLH: conceptualization, data curation, formal analysis, funding acquisition, investigation, methodology, project administration, resources, supervision, validation, visualization, writing – original draft, writing – reviewing and editing

## Data Availability

Data underlying all figures are included as individual data points within main and supplemental figures.

## Supplemental Material

**Supplemental Figure 1.**
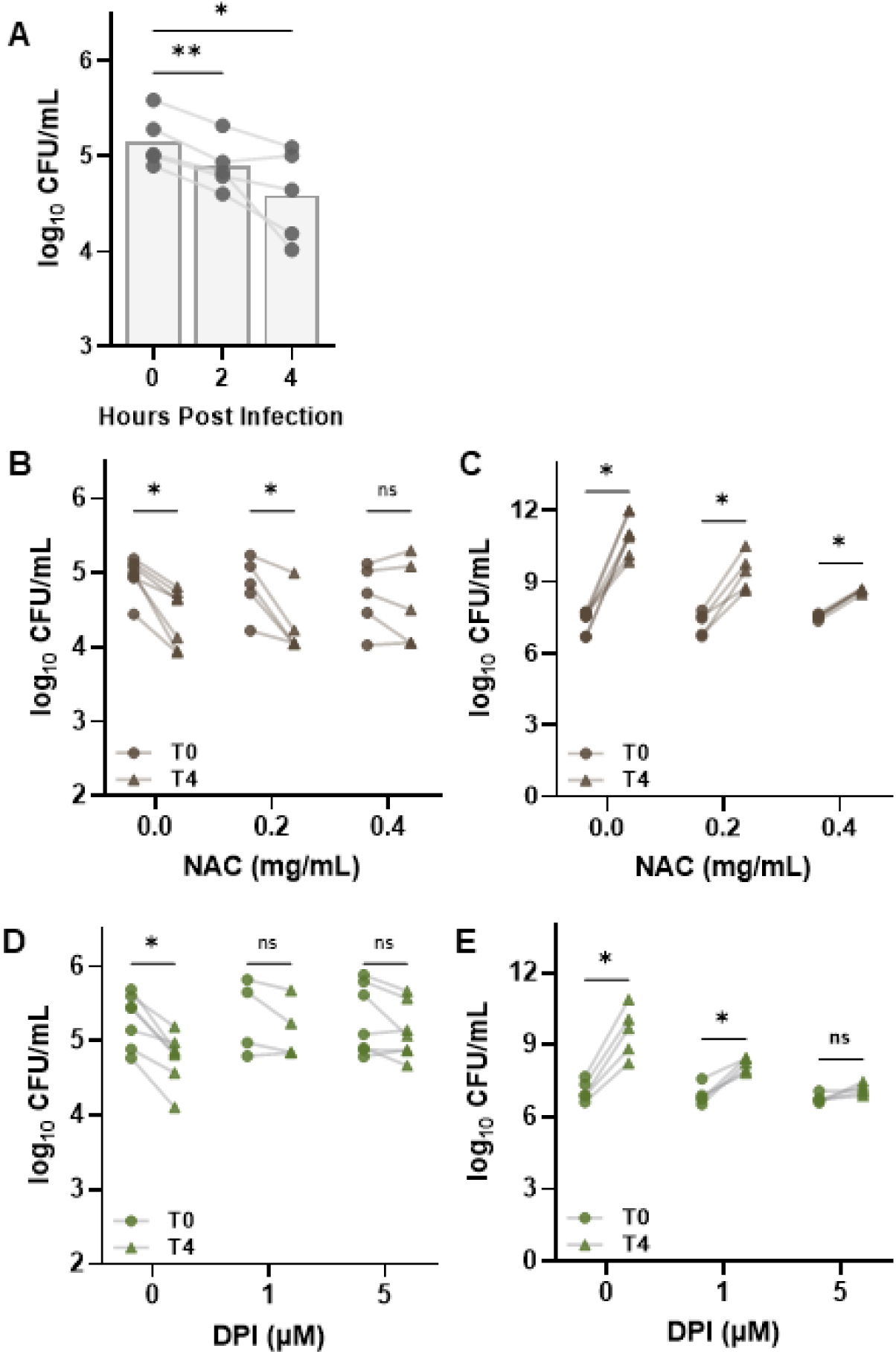
Chemical inhibition of reactive species increases intracellular *K. pneumoniae* survival. Macrophages (iBMDMs) were infected with KPPR1 for one hour prior to the removal of extracellular bacteria with gentamicin treatment. Abundance of intracellular bacteria was assessed immediately (T0), after two hours (T2), or after four hours (T4). In (A), the abundance of intracellular KPPR1 is displayed as log_10_ CFU/mL; n=4. \**p<*0.05, \*\**p<*0.01 by paired one-way ANOVA. (B-E) Chemical inhibitors of reactive species were assessed for their ability to influence macrophage killing of intracellular *K. pneumoniae* and affect overall bacterial viability. The top of the bar indicates the mean value. (B-C) The antioxidant N-acetylcysteine (NAC) was used to broadly inhibit reactive species at concentrations of 0.2mg/mL and 0.4mg/mL. (D-E) Diphenyleneiodonium (DPI) was used to inhibit NOX2 and iNOS activity at concentrations of 1µM and 5µM. In (B) and (D), iBMDMs were left untreated or treated with each inhibitor at the indicated concentration. Abundance of intracellular bacteria was assessed either immediately (T0) or after four hours (T4). In (C) and (E), the influence of each inhibitor on KPPR1 viability was assessed by incubating the bacteria with the inhibitors in culture media alone without macrophages. Bacterial abundance initially (T0) and after four hours (T4) was assessed by quantitative culture. In (B-E), \**p*<0.05 by paired *t-*test co paring log_10_ CFU/mL at T0 and T4 for each treatment; n=5-6 independent trials with symbols representing independent experiments.

**Supplemental Figure 2.**
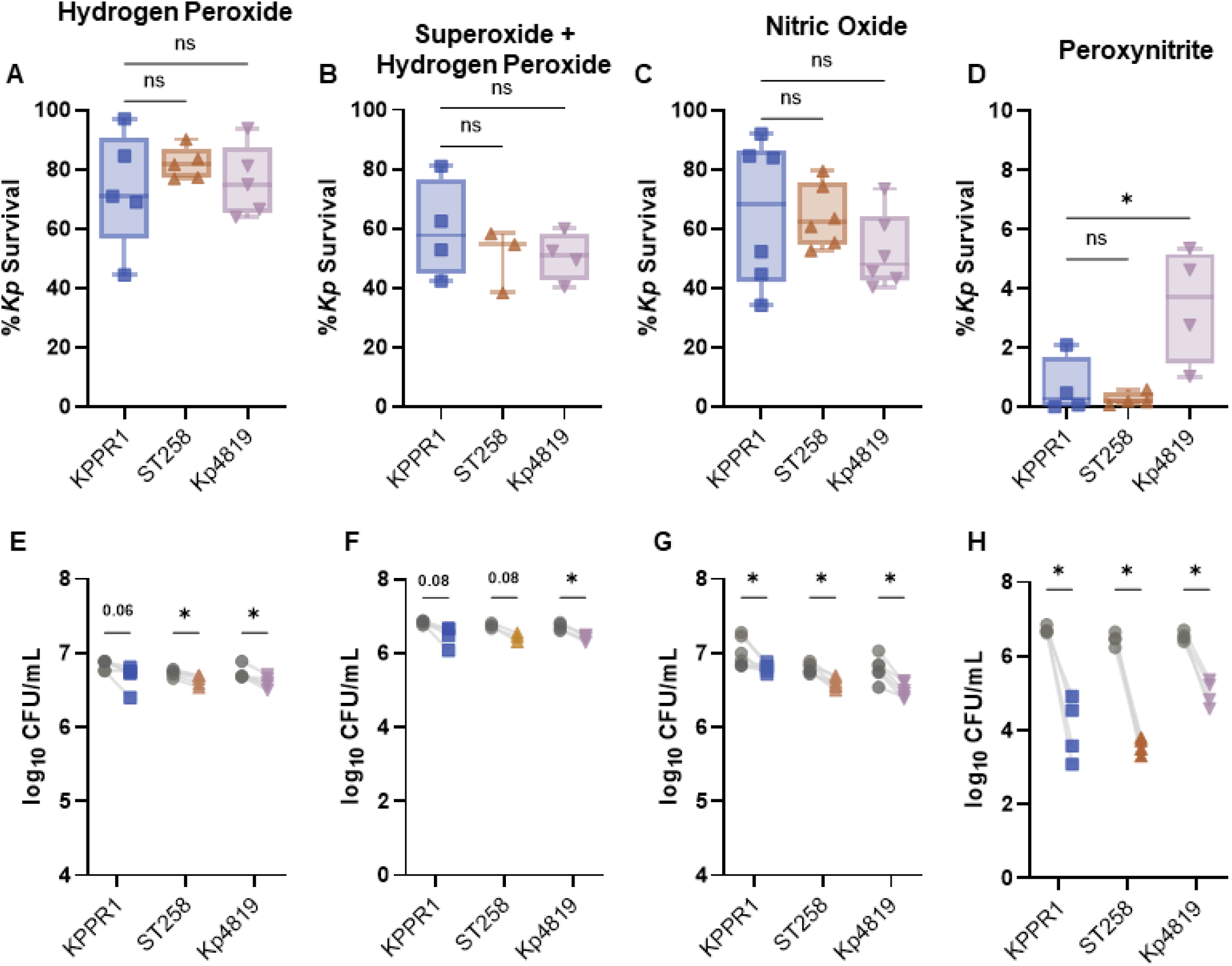
*K. pneumoniae* is susceptible to killing by diverse reactive species. The *K. pneumoniae* strains KPPR1, ST258, and Kp4819 were exposed to diverse reactive species and (A-D) percent survival or (E-H) log_10_ CFU/mL before and after exposure is displayed. Bacterial strains were exposed to (A, E) 1mM hydrogen peroxide, (B, F) 100 nU/mL xanthine + 10nM xanthine oxidase, (C, G) 2.5 mM DETA NONOate or (D, H) 1µM peroxynitrite. In (A-D), percent survival is calculated as (T0.5 or T2 CFU/T0 CFU)*100. In (A-D), \**p*<0.05 by one-way ANOVA comparing each value to KPPR1. In (E-H), \**p*<0.05 by paired *t-*test comparing CFU between each timepoin. Lines represent separate experiments; for all, n=3-6 independent trials.

**Supplemental Figure 3.**
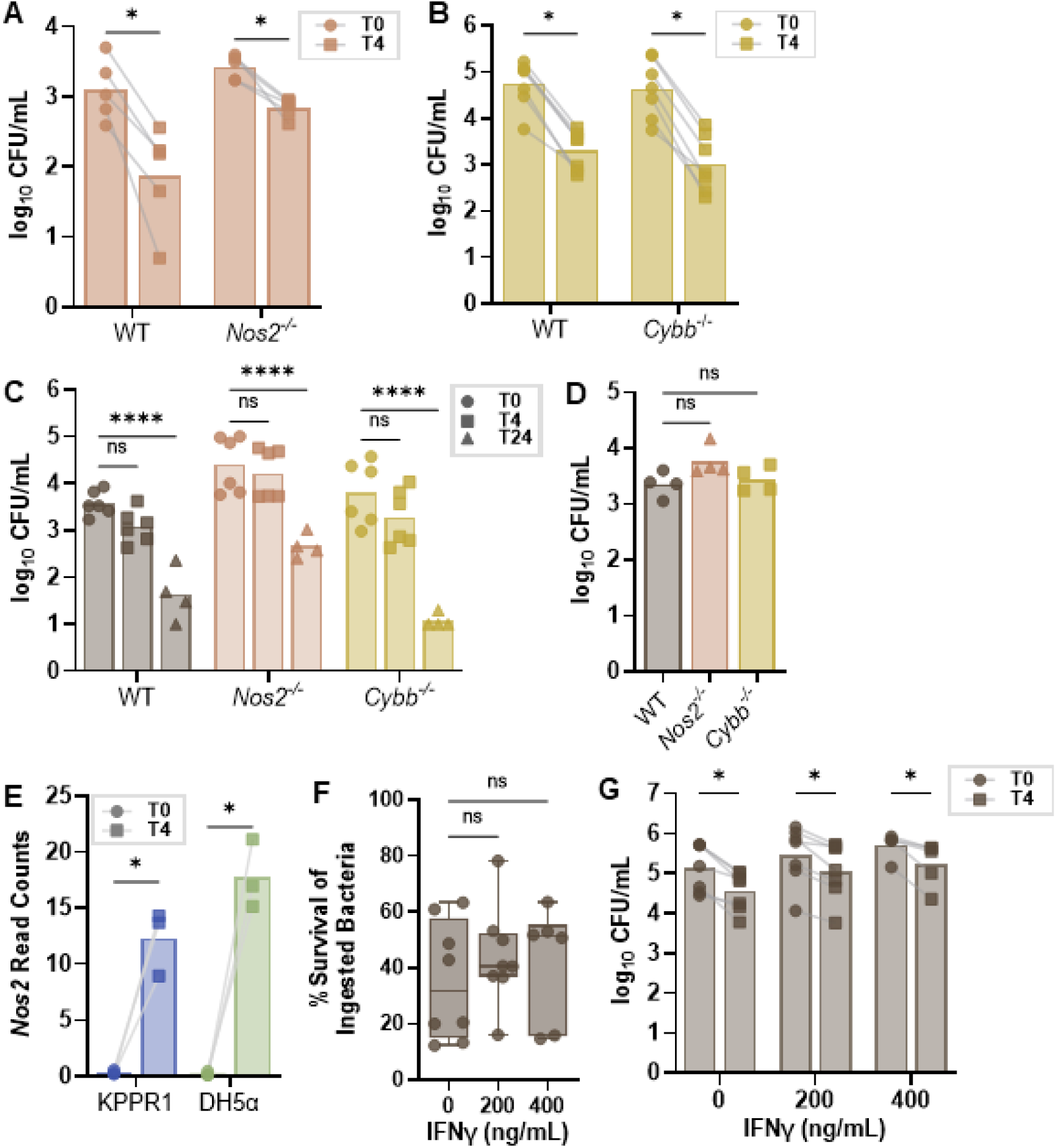
KPPR1 abundance recovered from monocyte-derived macrophages. The ability of wild-type, (A) *Nos2^-/-^*, or (B) *Cybb^-/-^*primary bone marrow-derived macrophages (pBMDMs) to kill intracellular KPPR1 was assessed using a gentamicin protection assay; n=5. (C) The results for WT, *Nos2^-/-^*, and *Cybb^-/-^* cells were validated using immortalized BMDMs. Abundance of intracellular bacteria was assessed either immediately (T0), after four hours (T4), or after 24 hours for panel C; n=4-6. (D) WT, *Nos2^-/-^*, and *Cybb^-/-^* iBMDMs were exposed to KPPR1 for 1 hour, treated with gentamicin, and immediately lysed to assess bacterial phagocytosis; n=4. (E) WT iBMDMs were infected with KPPR1 or *E. coli* (DH5α) and cellular RNA was collected at T0 and T4. *Nos2* counts per million are displayed; n=3. (F-G), WT, *Nos2^-/-^*, and *Cybb^-/-^* cells were left untreated (PBS) or stimulated with 200 or 400 ng/mL IFNγ for 16 hours prior to infection as described above. (F) Percent survival of ingested bacteria or (G) CFU/mL at T0 and T4 is displayed; n=5. In (A-B, E-G), \**p*<0.05 by paired *t-*test. In (C), \*\*\*\**p<*0.0001 by two-way ANOVA with Dunnett’s multiple correction. In (D), no differences between groups were detected by one-way ANOVA. For all, symbols represent independent experiments and the tops of bars indicate the mean value. For box plots in (F), box boundaries represent the 25^th^ and 75^th^ percentile ranges, the middle line represents the median value, and the whiskers represent the minimum and maximum values.

**Supplemental Figure 4.**
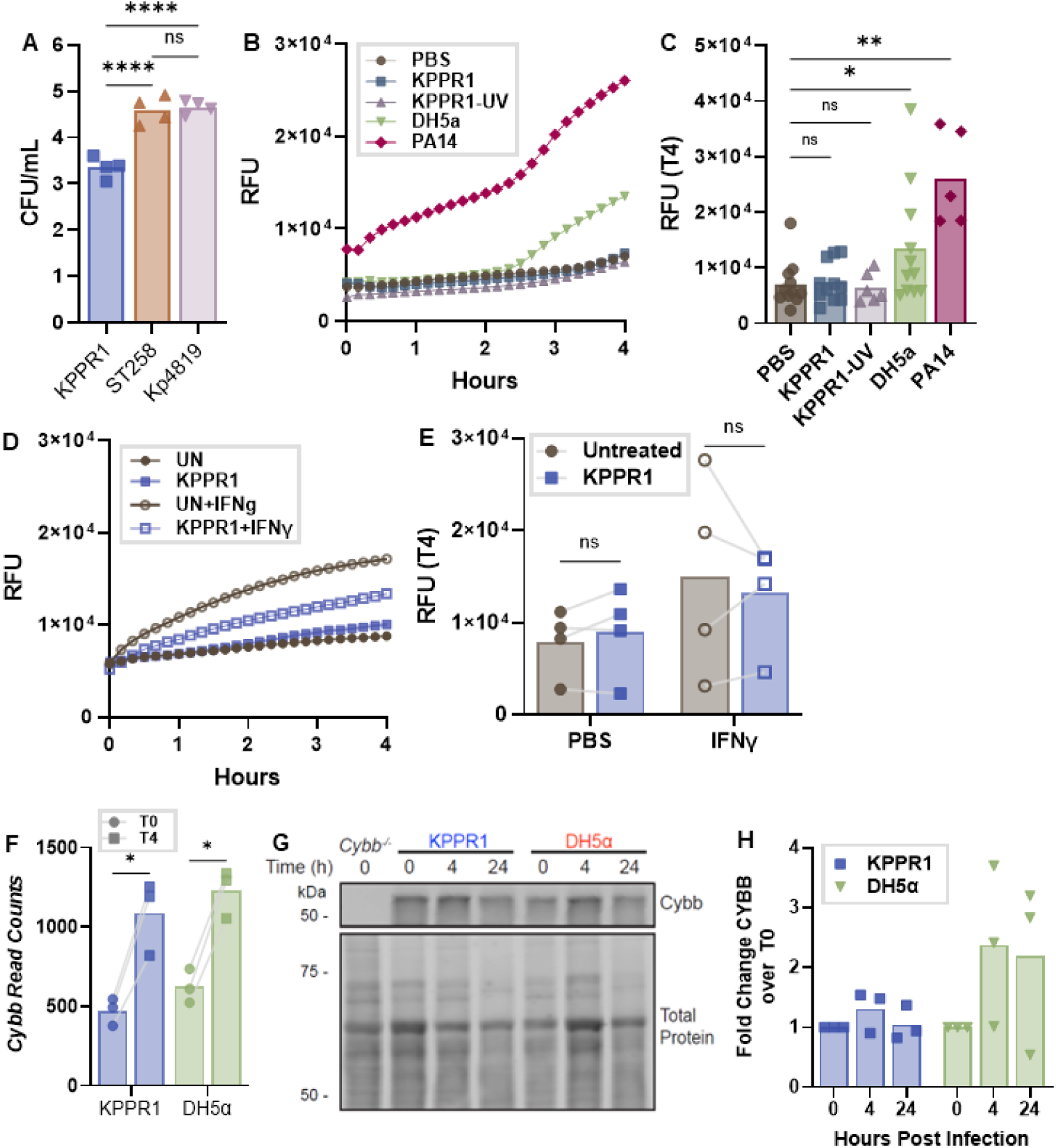
Macrophage ROS in response to infection from *K. pneumoniae* and other Gram-negative species. (A) WT immortalized bone marrow derived macrophages (iBMDMs) were assessed for the ability to phagocytize KPPR1, ST258, or Kp4819 by incubating the bacteria with cells for one hour, removing extracellular bacteria with gentamicin, and quantifying the abundance of intracellular bacteria; n=4. (B-C) iBMDMs were left untreated (PBS), or infected with *K. pneumoniae* (KPPR1), UV-killed KPPR1 (KPPR1-UV), *Escherichia coli* (DH5α), or *Pseudomonas aeruginosa* (PA14) for 1 hour. Cells were stained for ROS using DCFDA and fluorescence was measured every 10 minutes for 4 hours; n=5-11. (B) The mean DCFDA relative fluorescence (RFU) is displayed for each species. (C) The RFU at T4 for the data displayed in (B). (D-E) WT iBMDMs were left untreated (UN, PBS) or stimulated with 200ng/mL IFNγ prior to KPPR1 infection. DCFDA signaling was monitored as in (B-C); n=4. (F) WT iBMDMs were infected with KPPR1 or *E. coli* (DH5α) and cellular RNA collected at T0 and T4. *Cybb* counts per million are displayed; n=3. (G-H) WT iBMDMs were infected with KPPR1 or *E. coli* (DH5α) and total protein collected at T0, T4, and T24. Immunoblotting displays total protein and the signal from Cybb. (G) A representative blot from 3 independent trials is shown with *Cybb^-/-^* iBMDMs included as a negative control. (H) Densitometry of immunoblotting is: (Cybb signal/total protein signal) shown relative to the baseline signal. For (A, C), \*\**p*<0.01 and \*\*\*\**p*<0.0001 by one-way ANOVA. For (E, F), *\*p*<0.05 and \*\**p*<0.01 by paired *t-*test. For (H), no difference in Cybb signal over time was detected for either strain by two-way ANOVA. The top of each bar is the mean and symbols are independent trials.

**Supplemental Figure 5.**
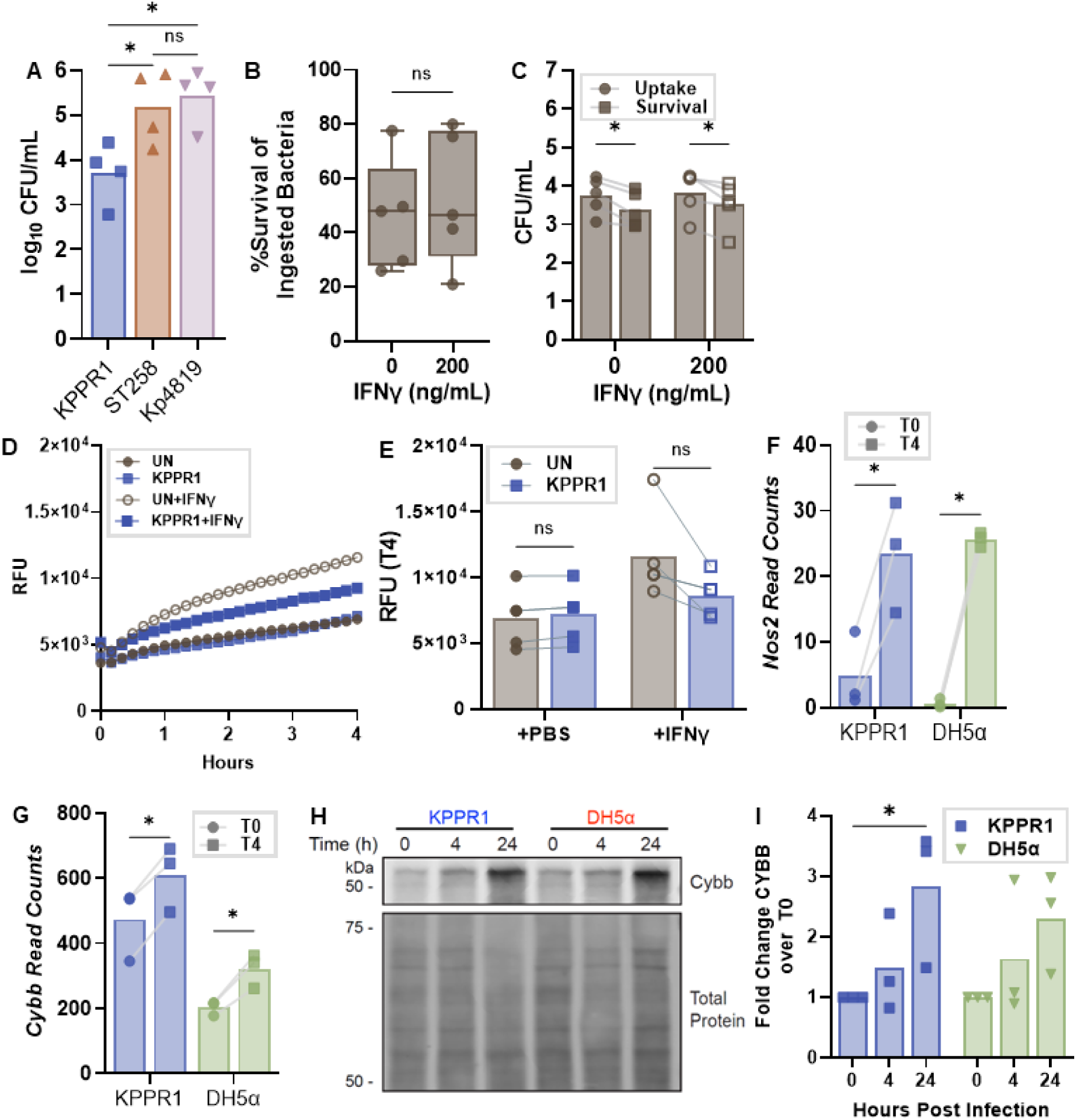
Alveolar-like macrophage ROS in response to *K. pneumoniae* infection. (A) Fetal liver-derived alveolar-like macrophages (FLAMs) were assessed for the ability to phagocytize KPPR1, ST258, or Kp4819 by incubating the bacteria with cells for one hour, removing extracellular bacteria with gentamicin, and quantifying the abundance of intracellular bacteria; n=4. (B-E) FLAMs were left untreated (PBS) or stimulated with 200ng/mL IFNγ prior to KPPR1 infection. In (B-C), FLAM bactericidal activity against *K. pneumoniae* was assessed using gentamicin protection assays. In (B), percent KPPR1 survival was assessed after 4 hours. In (C), the log_10_ CFU/mL at each timepoint in (B) is displayed; n=5. (D) FLAM ROS signal via DCFDA was monitored after KPPR1 infection every 10 minutes for 4 hours. The mean value is displayed; n=4. (E) The RFU at T4 for the data displayed in (D). (F-G) WT iBMDMs were infected with KPPR1 or *E. coli* (DH5α) and cellular RNA collected at T0 and T4. (F) *Nos2* or (G) *Cybb* counts per million are displayed; n=3. (H-I) FLAMs were infected with KPPR1 or *E. coli* (DH5α) and total protein collected at T0, T4, and T24. Immunoblotting displays total protein and the Cybb signal. (H) A representative blot independent trials. (I) Densitometry of immunoblotting is: (Cybb signal/total protein signal) from 3 shown relative to the baseline signal. \**p*<0.05 by two-way ANOVA. For (B-C, E-G), \**p*<0.05 by *t-*test. The top of each bar represents the mean, and symbols are independent trials.

## References

1. Singer M, Deutschman CS, Seymour CW, Shankar-Hari M, Annane D, Bauer M, Bellomo R, Bernard GR, Chiche J-D, Coopersmith CM, Hotchkiss RS, Levy MM, Marshall JC, Martin GS, Opal SM, Rubenfeld GD, van der Poll T, Vincent J-L, Angus DC. 2016. The Third International Consensus Definitions for Sepsis and Septic Shock (Sepsis-3). JAMA 315:801–810.

2. Hajj J, Blaine N, Salavaci J, Jacoby D. 2018. The “Centrality of Sepsis”: A Review on Incidence, Mortality, and Cost of Care. Healthcare (Basel) 6:90.

3. Novosad SA, Sapiano MRP, Grigg C, Lake J, Robyn M, Dumyati G, Felsen C, Blog D, Dufort E, Zansky S, Wiedeman K, Avery L, Dantes RB, Jernigan JA, Magill SS, Fiore A, Epstein L. 2016. Vital Signs: Epidemiology of Sepsis: Prevalence of Health Care Factors and Opportunities for Prevention. MMWR Morb Mortal Wkly Rep 65:864–869.

4. Diekema DJ, Hsueh P-R, Mendes RE, Pfaller MA, Rolston KV, Sader HS, Jones RN. 2019. The Microbiology of Bloodstream Infection: 20-Year Trends from the SENTRY Antimicrobial Surveillance Program. Antimicrob Agents Chemother 63:e00355–19.

5. Wisplinghoff H, Bischoff T, Tallent SM, Seifert H, Wenzel RP, Edmond MB. 2004. Nosocomial bloodstream infections in US hospitals: analysis of 24,179 cases from a prospective nationwide surveillance study. Clin Infect Dis 39:309–317.

6. Magill SS, Edwards JR, Bamberg W, Beldavs ZG, Dumyati G, Kainer MA, Lynfield R, Maloney M, McAllister-Hollod L, Nadle J, Ray SM, Thompson DL, Wilson LE, Fridkin SK, Emerging Infections Program Healthcare-Associated Infections and Antimicrobial Use Prevalence Survey Team. 2014. Multistate point-prevalence survey of health care-associated infections. N Engl J Med 370:1198–1208.

7. Holmes CL, Anderson MT, Mobley HLT, Bachman MA. 2021. Pathogenesis of Gram-Negative Bacteremia. Clin Microbiol Rev 34:e00234–20.

8. Vincent J-L, Rello J, Marshall J, Silva E, Anzueto A, Martin CD, Moreno R, Lipman J, Gomersall C, Sakr Y, Reinhart K, EPIC II Group of Investigators. 2009. International study of the prevalence and outcomes of infection in intensive care units. JAMA 302:2323–2329.

9. Holmes CL, Albin OR, Mobley HLT, Bachman MA. 2025. Bloodstream infections: mechanisms of pathogenesis and opportunities for intervention. Nat Rev Microbiol 23:210–224.

10. Antimicrobial Resistance Collaborators. 2022. Global burden of bacterial antimicrobial resistance in 2019: a systematic analysis. Lancet 399:629–655.

11. Sati H, Carrara E, Savoldi A, Hansen P, Garlasco J, Campagnaro E, Boccia S, Castillo-Polo JA, Magrini E, Garcia-Vello P, Wool E, Gigante V, Duffy E, Cassini A, Huttner B, Pardo PR, Naghavi M, Mirzayev F, Zignol M, Cameron A, Tacconelli E, WHO Bacterial Priority Pathogens List Advisory Group. 2025. The WHO Bacterial Priority Pathogens List 2024: a prioritisation study to guide research, development, and public health strategies against antimicrobial resistance. Lancet Infect Dis 25:1033–1043.

12. Holmes CL, Smith SN, Gurczynski SJ, Severin GB, Unverdorben LV, Vornhagen J, Mobley HLT, Bachman MA. 2022. The ADP-Heptose Biosynthesis Enzyme GmhB is a Conserved Gram-Negative Bacteremia Fitness Factor. Infect Immun 90:e0022422.

13. Wong Fok Lung T, Charytonowicz D, Beaumont KG, Shah SS, Sridhar SH, Gorrie CL, Mu A, Hofstaedter CE, Varisco D, McConville TH, Drikic M, Fowler B, Urso A, Shi W, Fucich D, Annavajhala MK, Khan IN, Oussenko I, Francoeur N, Smith ML, Stockwell BR, Lewis IA, Hachani A, Upadhyay Baskota S, Uhlemann A-C, Ahn D, Ernst RK, Howden BP, Sebra R, Prince A. 2022. Klebsiella pneumoniae induces host metabolic stress that promotes tolerance to pulmonary infection. Cell Metab 34:761–774.e9.

14. Xiong H, Carter RA, Leiner IM, Tang Y-W, Chen L, Kreiswirth BN, Pamer EG. 2015. Distinct Contributions of Neutrophils and CCR2+ Monocytes to Pulmonary Clearance of Different Klebsiella pneumoniae Strains. Infect Immun 83:3418–3427.

15. Chen L, Zhang Z, Barletta KE, Burdick MD, Mehrad B. 2013. Heterogeneity of lung mononuclear phagocytes during pneumonia: contribution of chemokine receptors. Am J Physiol Lung Cell Mol Physiol 305:L702–711.

16. Broug-Holub E, Toews GB, van Iwaarden JF, Strieter RM, Kunkel SL, Paine R, Standiford TJ. 1997. Alveolar macrophages are required for protective pulmonary defenses in murine Klebsiella pneumonia: elimination of alveolar macrophages increases neutrophil recruitment but decreases bacterial clearance and survival. Infect Immun 65:1139–1146.

17. Vazquez-Torres A, Jones-Carson J, Mastroeni P, Ischiropoulos H, Fang FC. 2000. Antimicrobial actions of the NADPH phagocyte oxidase and inducible nitric oxide synthase in experimental salmonellosis. I. Effects on microbial killing by activated peritoneal macrophages in vitro. J Exp Med 192:227–236.

18. Nguyen JA, Yates RM. 2021. Better Together: Current Insights Into Phagosome-Lysosome Fusion. Front Immunol 12:636078.

19. Fang FC. 2004. Antimicrobial reactive oxygen and nitrogen species: concepts and controversies. Nat Rev Microbiol 2:820–832.

20. Wilcox AE, Andres CJ, Bachman MA, Holmes CL. 2026. Klebsiella pneumoniae factors enhancing bacteremia have distinct contributions to macrophage-mediated, oxidative, and nitrosative stress resistance. Infect Immun 94:e0073925.

21. Hickman-Davis JM, O’Reilly P, Davis IC, Peti-Peterdi J, Davis G, Young KR, Devlin RB, Matalon S. 2002. Killing of Klebsiella pneumoniae by human alveolar macrophages. Am J Physiol Lung Cell Mol Physiol 282:L944–956.

22. Hickman-Davis J, Gibbs-Erwin J, Lindsey JR, Matalon S. 1999. Surfactant protein A mediates mycoplasmacidal activity of alveolar macrophages by production of peroxynitrite. Proc Natl Acad Sci U S A 96:4953–4958.

23. Gillissen A, Nowak D. 1998. Characterization of N-acetylcysteine and ambroxol in anti-oxidant therapy. Respir Med 92:609–623.

24. Hampton MB, Winterbourn CC. 1995. Modification of neutrophil oxidant production with diphenyleneiodonium and its effect on bacterial killing. Free Radic Biol Med 18:633–639.

25. Stuehr DJ, Fasehun OA, Kwon NS, Gross SS, Gonzalez JA, Levi R, Nathan CF. 1991. Inhibition of macrophage and endothelial cell nitric oxide synthase by diphenyleneiodonium and its analogs. FASEB J 5:98–103.

26. Werner JL, Escolero SG, Hewlett JT, Mak TN, Williams BP, Eishi Y, Núñez G. 2017. Induction of Pulmonary Granuloma Formation by Propionibacterium acnes Is Regulated by MyD88 and Nox2. Am J Respir Cell Mol Biol 56:121–130.

27. Russo TA, Alvarado CL, Davies CJ, Drayer ZJ, Carlino-MacDonald U, Hutson A, Luo TL, Martin MJ, Corey BW, Moser KA, Rasheed JK, Halpin AL, McGann PT, Lebreton F. 2024. Differentiation of hypervirulent and classical Klebsiella pneumoniae with acquired drug resistance. mBio 15:e0286723.

28. Snitkin ES, Zelazny AM, Thomas PJ, Stock F, NISC Comparative Sequencing Program Group, Henderson DK, Palmore TN, Segre JA. 2012. Tracking a hospital outbreak of carbapenem-resistant Klebsiella pneumoniae with whole-genome sequencing. Sci Transl Med 4:148ra116.

29. Unverdorben LV, Pirani A, Gontjes K, Moricz B, Holmes CL, Snitkin ES, Bachman MA. 2025. Klebsiella pneumoniae evolution in the gut leads to spontaneous capsule loss and decreased virulence potential. mBio 16:e0236224.

30. Managò A, Becker KA, Carpinteiro A, Wilker B, Soddemann M, Seitz AP, Edwards MJ, Grassmé H, Szabò I, Gulbins E. 2015. Pseudomonas aeruginosa pyocyanin induces neutrophil death via mitochondrial reactive oxygen species and mitochondrial acid sphingomyelinase. Antioxid Redox Signal 22:1097–1110.

31. Karges J. 2026. Reactive Oxygen Species Detection with Fluorescent Probes: Limitations and Recommendations beyond DCFH-DA. J Med Chem 69:1970–1981.

32. Traore K, Trush MA, George M, Spannhake EW, Anderson W, Asseffa A. 2005. Signal transduction of phorbol 12-myristate 13-acetate (PMA)-induced growth inhibition of human monocytic leukemia THP-1 cells is reactive oxygen dependent. Leuk Res 29:863–879.

33. Fontayne A, Dang PM-C, Gougerot-Pocidalo M-A, El-Benna J. 2002. Phosphorylation of p47phox sites by PKC alpha, beta II, delta, and zeta: effect on binding to p22phox and on NADPH oxidase activation. Biochemistry 41:7743–7750.

34. Thomas ST, Wierenga KA, Pestka JJ, Olive AJ. 2022. Fetal Liver-Derived Alveolar-like Macrophages: A Self-Replicating Ex Vivo Model of Alveolar Macrophages for Functional Genetic Studies. Immunohorizons 6:156–169.

35. Guilliams M, Scott CL. 2017. Does niche competition determine the origin of tissue-resident macrophages? Nat Rev Immunol 17:451–460.

36. Holmes CL, Wilcox AE, Forsyth V, Smith SN, Moricz BS, Unverdorben LV, Mason S, Wu W, Zhao L, Mobley HLT, Bachman MA. 2023. Klebsiella pneumoniae causes bacteremia using factors that mediate tissue-specific fitness and resistance to oxidative stress. PLoS Pathog 19:e1011233.

37. Holmes CL, Dailey KG, Hullahalli K, Wilcox AE, Mason S, Moricz BS, Unverdorben LV, Balazs GI, Waldor MK, Bachman MA. 2025. Patterns of Klebsiella pneumoniae bacteremic dissemination from the lung. Nat Commun 16:785.

38. Nguyen GT, Shaban L, Mack M, Swanson KD, Bunnell SC, Sykes DB, Mecsas J. 2020. SKAP2 is required for defense against K. pneumoniae infection and neutrophil respiratory burst. Elife 9:e56656.

39. Tsai WC, Strieter RM, Zisman DA, Wilkowski JM, Bucknell KA, Chen GH, Standiford TJ. 1997. Nitric oxide is required for effective innate immunity against Klebsiella pneumoniae. Infect Immun 65:1870–1875.

40. Lu G, Zhang R, Geng S, Peng L, Jayaraman P, Chen C, Xu F, Yang J, Li Q, Zheng H, Shen K, Wang J, Liu X, Wang W, Zheng Z, Qi C-F, Si C, He JC, Liu K, Lira SA, Sikora AG, Li L, Xiong H. 2015. Myeloid cell-derived inducible nitric oxide synthase suppresses M1 macrophage polarization. Nat Commun 6:6676.

41. Nguyen HK, Duke MM, Grayton QE, Broberg CA, Schoenfisch MH. 2024. Impact of nitric oxide donors on capsule, biofilm and resistance profiles of Klebsiella pneumoniae. Int J Antimicrob Agents 64:107339.

42. Paczosa MK, Silver RJ, McCabe AL, Tai AK, McLeish CH, Lazinski DW, Mecsas J. 2020. Transposon Mutagenesis Screen of Klebsiella pneumoniae Identifies Multiple Genes Important for Resisting Antimicrobial Activities of Neutrophils in Mice. Infect Immun 88:e00034–20.

43. Wyres KL, Lam MMC, Holt KE. 2020. Population genomics of Klebsiella pneumoniae. Nat Rev Microbiol 18:344–359.

44. Paczosa MK, Mecsas J. 2016. Klebsiella pneumoniae: Going on the Offense with a Strong Defense. Microbiol Mol Biol Rev 80:629–661.

45. Xiong H, Keith JW, Samilo DW, Carter RA, Leiner IM, Pamer EG. 2016. Innate Lymphocyte/Ly6C(hi) Monocyte Crosstalk Promotes Klebsiella Pneumoniae Clearance. Cell 165:679–689.

46. Holmes CL. 2023. mSphere of Influence: Coordinating effective clearance of bacterial bloodstream infections. mSphere 8:e0052123.

47. Broberg CA, Wu W, Cavalcoli JD, Miller VL, Bachman MA. 2014. Complete Genome Sequence of Klebsiella pneumoniae Strain ATCC 43816 KPPR1, a Rifampin-Resistant Mutant Commonly Used in Animal, Genetic, and Molecular Biology Studies. Genome Announc 2:e00924–14.

48. Rao K, Patel A, Sun Y, Vornhagen J, Motyka J, Collingwood A, Teodorescu A, Baang JH, Zhao L, Kaye KS, Bachman MA. 2021. Risk Factors for Klebsiella Infections among Hospitalized Patients with Preexisting Colonization. mSphere 6:e0013221.

49. Kiritsy MC, Ankley LM, Trombley J, Huizinga GP, Lord AE, Orning P, Elling R, Fitzgerald KA, Olive AJ. 2021. A genetic screen in macrophages identifies new regulators of IFNγ-inducible MHCII that contribute to T cell activation. Elife 10:e65110.

50. Gilliland HN, Beckman OK, Olive AJ. 2023. A Genome-Wide Screen in Macrophages Defines Host Genes Regulating the Uptake of Mycobacterium abscessus. mSphere 8:e0066322.

51. Fejer G, Wegner MD, Györy I, Cohen I, Engelhard P, Voronov E, Manke T, Ruzsics Z, Dölken L, Prazeres da Costa O, Branzk N, Huber M, Prasse A, Schneider R, Apte RN, Galanos C, Freudenberg MA. 2013. Nontransformed, GM-CSF-dependent macrophage lines are a unique model to study tissue macrophage functions. Proc Natl Acad Sci U S A 110:E2191–2198.

52. Brian BF, Guerrero CR, Freedman TS. 2020. Immunopharmacology and Quantitative Analysis of Tyrosine Kinase Signaling. Curr Protoc Immunol 130:e104.

